# ST-FFPE-mIF: Integrating Spatial Transcriptomics and Multiplex Immunofluorescence in Formalin-Fixed Paraffin-Embedded Tissues Using Stereo-seq

**DOI:** 10.1101/2025.01.24.634655

**Authors:** Xue Zhang, Meng Zhang, Yuan Xu, Yangyang Song, Xinfeng Yang, Yanan Wu, Xuelin Zhao, Dongsheng Ran, Xin Liu, Huaqiang Huang, Wenxiao Lei, Hongyan Li, Yongfen Zhang, Feng Xi, Guibo Li, Xing Liu, Luohao Xu, Ao Chen, Sha Liao, Jiajun Zhang

## Abstract

Stereo-seq is the state-of-the-art technique, which allows capturing of spatial transcriptomic data at sub-cellular resolution and in large scale of view. Stereo-seq FFPE technique is recently launched with the capability to obtain spatially resolved gene expression in FFPE tissue sections. However, the omic information is limited to transcriptome. To delineate spatial molecular hierarchy from transcriptome to phenome, we present the protocol that implements nine-plex immunofluorescence staining with detection of spatial transcriptomics in one FFPE tissue section (short for ST-FFPE-mIF). This protocol details the modified biochemistry procedure and also provides analytical framework to achieve single-cell resolution through nuclear or membrane staining-based cell segmentation. We demonstrated the application of ST-FFPE-mIF in colorectal cancer sample to illustrate how ST-FFPE-mIF facilitates a robust analysis of spatially resolved multi-omic data at the single-cell level.

## Introduction

Spatial transcriptomics (ST) technology is a cutting-edge biological technology enabling transcriptomic profiling within a spatial context to revolutionize biological research[1–3]. The commercial ST technologies can be classified into imaging-based such as MERFISH, seqFISH, and osmFISH, and sequencing-based methods including 10X Visium, Slide-seq V2, and Stereo-seq[2, 4, 5]. Despite the high resolution, imaging-based ST methods present several notable disadvantages: low throughput, complexity, high costs, time consumption, and comparability of transcripts[6]. Although addressing the challenges of low throughput[7], most sequencing-based ST methods are limited by resolution constraints. Emerging of advanced ST technology, like

Stereo-seq, overcomes the mentioned limitation. Stereo-seq utilizes a DNA nanoball (DNB) patterned chip with a center-to-center distance of 500 nm, enabling the detection of spatially resolved and unbiased transcriptomic profiling at subcellular resolution[8]. A newly released version of Stereo-seq upgraded the probes with randomers (Stereo-seq-FFPE), which made the technique compatible with the formalin-fixed, paraffin-embedded (FFPE) sample.

ST technology allows for the visualization and analysis of gene expression patterns within the spatial context of tissue sections. However, unimodal (transcriptome) measurement also limits the application to link the mRNA information to functional activities. Given that protein markers serve as the gold standard for indicating biological functions and identifying specific cell types, it is necessary to develop a technique to add protein expression on top of ST data for the multi-modal analysis. The currently available spatial co-detection technologies exhibit several limitations, including a lack of sub-cellular resolution in Visium, DBiT-seq and SM-Omics[6], unable to achieve unbiased transcriptomic profiling in Merfish[9], and the necessity for costly equipment in DSP[10]and Visium HD[11]. These challenges hinder the potential of these spatial techniques to adequately fulfill the requirements of clinical applications and translational research[12, 13]. The microscope-based multiplexing technique represents a pathology-related and relatively time-efficient method for assessing protein expression[14]. In light of the aforementioned, there is a clear and pressing need to develop a multi-omics co-detection technology with subcellular resolution capable of performing spatial whole-transcriptome capturing and multiplex immunofluorescence staining.

This study presents an innovative protocol termed Spatial Transcriptome and Multiplex Immunofluorescence for Paraffin-Embedded Samples (ST-FFPE-mIF). This methodology facilitates the simultaneous detection of nine-plex immunofluorescence staining in conjunction with whole transcriptome by Stereo-seq on one section of FFPE tissue. The protocol of ST-FFPE-mIF needs less than four hours for mIF staining process. Furthermore, the solution includes a comprehensive analytical workflow for image processing, data visualization, and multi-omic data integration. ST-FFPE-mIF also provides a recommended guideline for cell segmentation based on either nuclear or membrane staining. Additionally, we demonstrate the application of ST-FFPE-mIF in investigating cellular and molecular landscape in colorectal cancer (CRC) by integrated analysis of spatial transcriptomic data and mIF images at single-cell resolution. This innovative solution has the potential to enhance the understanding of physiological and pathological processes through the integration of multi-omic and architectural data.

## Results

### Workflow of ST-FFPE-mIF on Formalin-fixed paraffin-embedded sample

The Stereo-seq-FFPE technology is a powerful method that enabling precise and accurate visualization of spatial mRNA information at single-cell resolution in FFPE tissue. We have enhanced the capabilities of Stereo-seq-FFPE by providing a protocol (Supplementary 1) to enable the mIF staining before the standard ST protocol, referred to as ST-FFPE-mIF(Fig. 1, step 1). To overcome the challenges associated with significant crosslinking in FFPE samples, we implemented a refined step for antigen retrieval (Supplementary 1) to facilitate the sufficient unmasking of antigenic epitopes while preventing the degradation of RNA molecules (Fig. 1, step 2). Given the limited antibody sources, fluorescence-labeled conjugated (FL-conjugated) primary antibodies were utilized for mIF staining to ensure compatibility with various different microscope systems[15]. To achieve the maximum number of targets for mIF, a protocol was developed for the elution of antibodies from the tissue section after 1^st^ round imaging, which enabled a subsequent 2^nd^ round of mIF staining and imaging (Fig 1, step 2).

**Fig 1.**
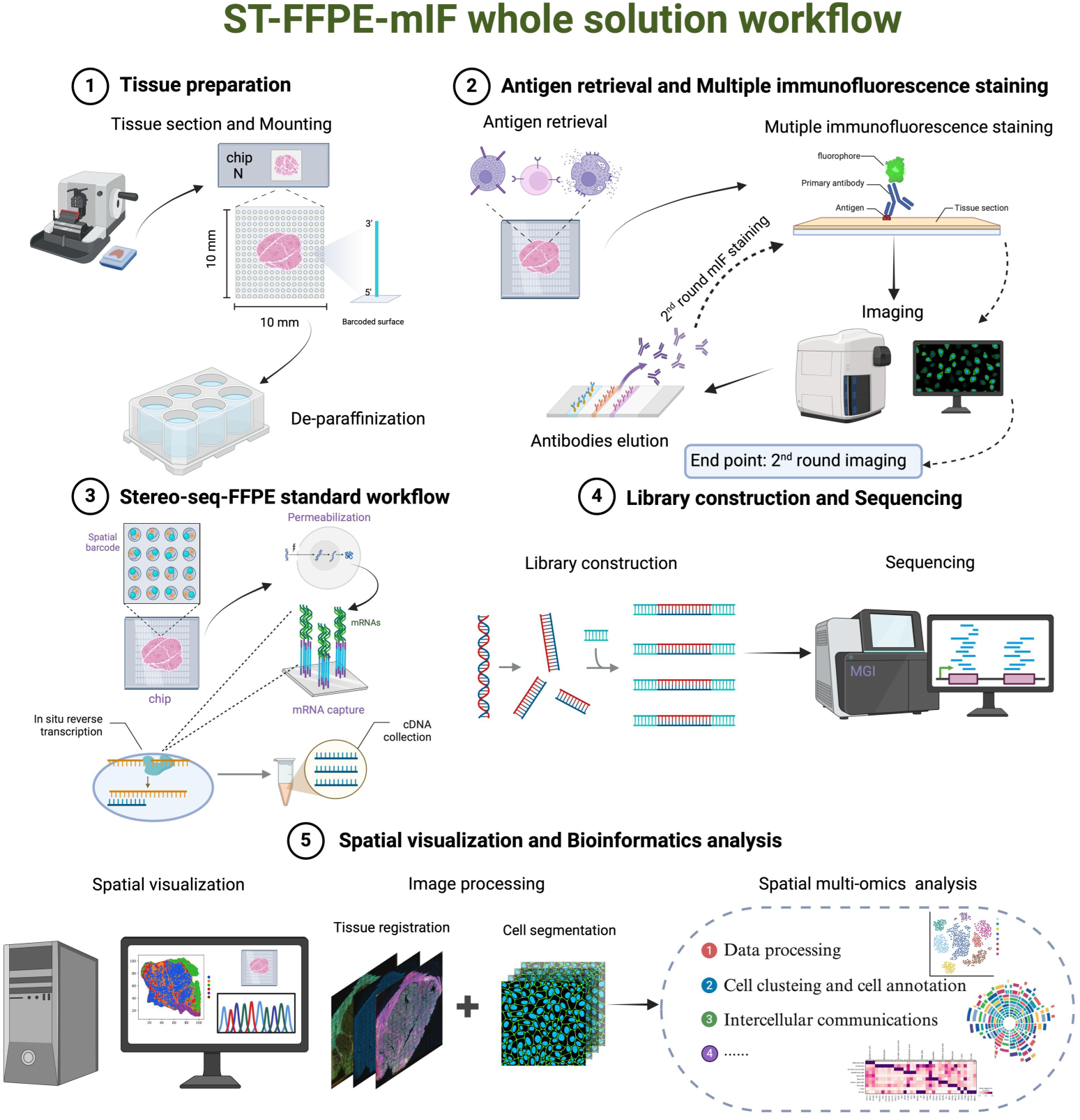
Schematic workflow and bioinformatics analysis of the ST-FFPE-mIF whole solution. (1) FFPE tissue section was mounted on the chip and performed de-paraffinization. (2) The section on the chip was subjected to antigen repair, blocking, antibody incubation, and mIF imaging. Before the second round of mIF staining, antibodies from the 1^st^ round should be removal, and the epitope would be re-exposed. Then, the above mIF protocol should be repeated to generate another mIF image. Nine-colored mIF images were obtained accordingly. (3) In situ RNA capture and cDNA synthesis were conducted after permeabilization. (4) cDNA amplification, library construction, and sequencing were performed according to manufacturer’s protocol to obtain Stereo-seq sequencing result. (5) mIF images and spatially resolved gene expression was subjected to downstream spatial visualization and bioinformatics analyses including image processing and spatial multi-omics analysis to address scientific questions.

After the 2^nd^ round of mIF imaging, the protocol of Stereo-seq-FFPE should be followed, including permeabilization, probe hybridization, in situ reverse transcription (Fig. 1, step 3), library construction and sequencing (Fig. 1, step 4). Following rigorous data processing, a spatially resolved gene expression matrix is generated that can be aligned with mIF images. The aligned images are subjected to cell segmentation, yielding a single-cell resolved expression matrix that enables advanced spatial multi-omics analyses (Fig. 1, step 5). A detailed workflow of the ST-FFPE-mIF and a step-by-step protocol can be consulted in the supplementary section.

### Evaluation of spatially resolved transcriptomic profiles and mIF images from ST-FFPE-mIF in the human colorectal cancer FFPE sample

To evaluate the performance of ST-FFPE-mlF, we applied it to FFPE sections of human CRC. The tested mIF panel was designed to identify cancerous regions (HER2, Villin, P53, CK19), T cell (CD3), macrophage (CD68), and fibroblast (SMA, Vimentin). ST-FFPE-mIF detected an average of 798 genes and 1235 unique molecular identifiers (UMIs) per bin50 (25 µm × 25 µm). Both the mIF staining image and spatial distribution heat maps of detected gene types and UMI counts displayed a strong concordance with histomorphology (H&E Staining) (Fig. 2A). The detailed mIF images accurately reflected the histological structure, and the IF images stained by each marker had its unique spatial distributions and specificities (Fig. 2B).

**Fig 2.**
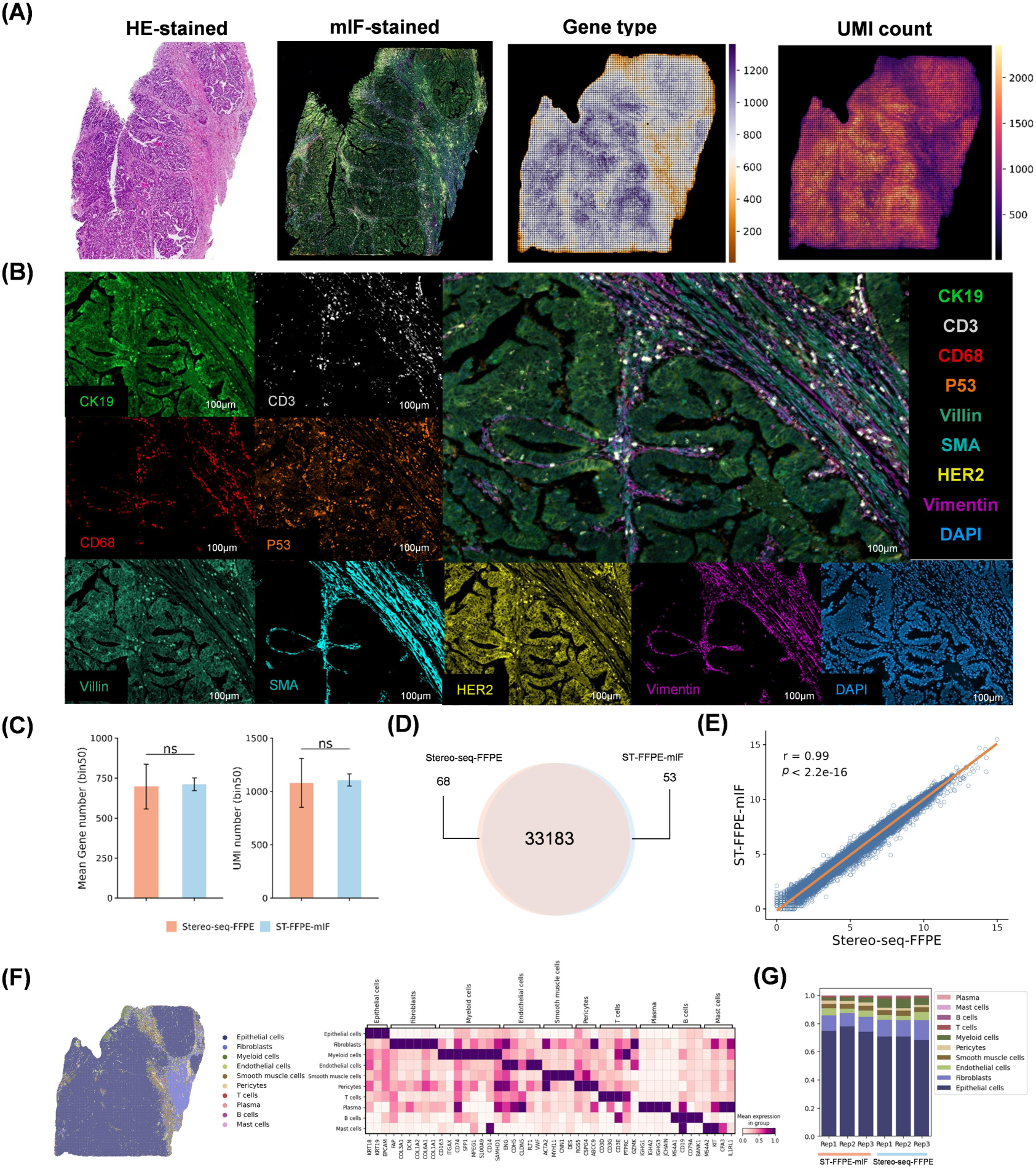
Evaluation of performance of ST-FFPE-mIF using human colorectal cancer section. (A) ST-FFPE-mIF profiling of a human colorectal cancer FFPE section. Left: H&E staining of an adjacent section. Middle left: Co-detection of mIF image. Middle right: Spatial gene expression count map at bin50. Right: Spatial UMI count map at bin50. (B) Representative images for mIF staining of 8-plex antibody panel in CRC tissue. Cells were colored by CK19 (green), CD3 (gray), CD68 (red), P53 (orange), Villin (blue green), SMA (turquoise), HER2 (yellow), Vimentin (violet), DAPI (blue). Scale bars = 100 µm. (C) Bar plots show the distribution of unique genes and UMI counts for Stereo-seq-FFPE (n = 3) and ST-FFPE-mIF (n = 3) data generated from consecutive tissue sections from the same CRC tissue specimen. (D) The Venn diagram shows the confluence and variance in gene capture between Stereo-seq-FFPE and ST-FFPE-mIF. (E) Scatter plot shows the gene expression significantly correlated between Stereo-seq-FFPE and ST-FFPE-mIF. Correlation coefficients were determined by the Pearson correlation coefficient. (F) Cell annotation is performed based on transcriptomic data at bin50 resolution. Representative figures show spatial visualization of cell identification and heatmap depicting the expression of classic markers of 10 annotated cell types. (G) Bar chart displays the percentage of each cell type within the analyzed sample.

Comparative assessment between results from ST-FFPE-mIF and standard Stereo-seq-FFPE was performed to estimate the capture efficiency. The sequencing depth was meticulously controlled, employing triplicate continuous sections for each methodology as statistical replicates. Figure 2C indicated that ST-FFPE-mIF could profile the spatial transcriptome with a comparable number of detected genes and UMIs per spot when compared to standard Stereo-seq-FFPE. Moreover, there was a significant overlap of detected gene types (Fig. 2D) and a strong concordance in gene expression (Fig. 2E) between the two techniques.

In addition to evaluating general capture efficiency, we further compare the annotation results. We annotated the ST data at a resolution of bin50 using cell2location[16, 17], in conjunction with a corresponding single-cell RNA sequence (scRNA-seq) reference[18]. The distribution of annotated cell types was consistent with the morphological features revealed by H&E staining of the adjacent section. Marker genes of the corresponding cell types expressed specially, demonstrating the accuracy of annotation (Fig. 2F, Fig. S1, and Fig S2A). We also observed a strong concordance in the proportions of annotated cell types across the two techniques (Fig. 2G). This suggests that the spatial transcriptomic data acquired through the ST-FFPE-mIF maintains a high consistency with the data generated by the commercially available Stereo-seq-FFPE technology. Collectively, these results indicate that ST-FFPE-mIF possessed the ability to obtain multi-omics data without compromising the capture efficiency.

### ST-FFPE-mIF guarantees the production of authentic, high-quality mIF images

Chromogenic immunohistochemistry (IHC) is widely employed to assess protein biomarker expression in FFPE tissue, which is widely used in pathology[19]. To verify the precision and reliability of mIF staining in ST-FFPE-mIF, we utilized adjacent sections for IHC staining of each target within the eight-plex panel. This approach facilitated a direct comparison of signal consistency within identical fields of view (FOV), thereby enabling an unequivocal assessment of individual IF results in relation to IHC outcomes from corresponding regions. Despite the unavoidable effects of intrinsic autofluorescence and signal channel crosstalk[20, 21], the tissue-specific structural features and positive signals observed between IHC and individual IF demonstrated a high degree of concordance (Fig. 3A). Furthermore, we employed ImageJ software to calculate the average area of positive cells across 20 randomly selected FOVs in each image, which permitted a comparative analysis of IHC and IF results based on specific targets. The comparison revealed no significant differences in positive areas between mIF and IHC for each target (Fig. 3B). It is important to note that discrepancies in positive areas may stem from variations in imaging principles and detection sensitivities to IHC versus IF methodologies. In summary, our mIF images exhibited authentic and reliable fluorescence signals that could support precise identification of cells types within tumor microenvironment.

**Fig 3.**
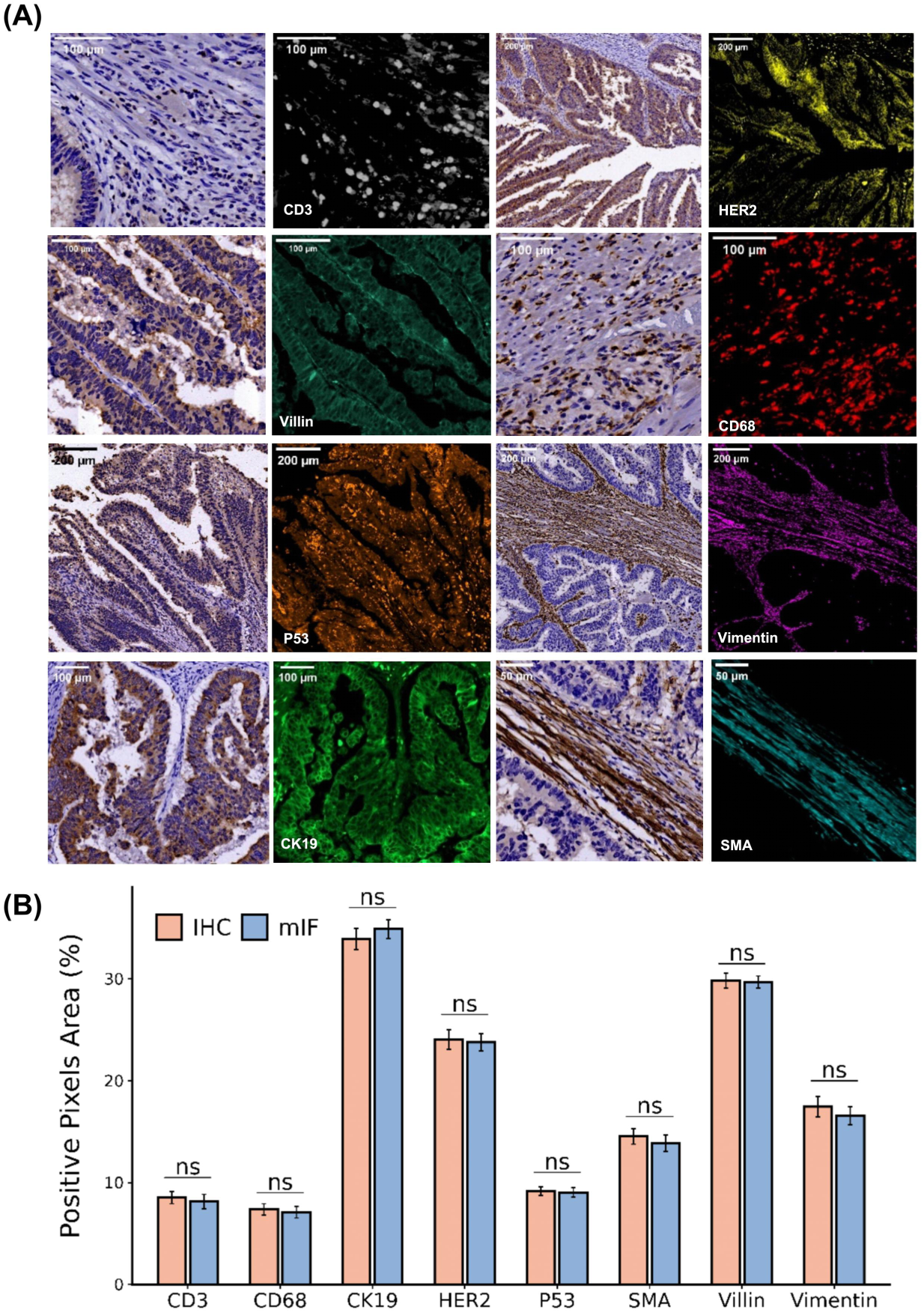
Assess the accuracy of mIF staining from ST-FFPE-mIF staining by using chromogenic IHC as gold standard. (A) Representative FOVs in photomicrograph of mIF and chromogenic IHC show comparable results. (B) Quantitative assessment of positive signal in each image demonstrates equivalence of staining between mIF and IHC. ns: not significant.

### Evaluation of technical reproducibility of ST-FFPE-mIF

Three adjacent sections from the same human CRC sample were utilized to assess technical reproducibility. The ST-FFPE-mIF datasets generated from these samples revealed an outstanding consistency across several fundamental metrics at bin50 resolution. This included the number of genes and UMIs that exhibited similar distribution in violin plots, a high overlap of captured gene types, and correlations of captured gene profiles exceeding 0.99 (Fig. 4A-C). The spatial distributions, compositional proportions, and expression levels of classic marker genes in the corresponding cell types of annotated cells across the three replicates were similar (Fig. 4D and Fig. S2A). Furthermore, the gene profiles among the individual cell types also showed high correlations (Fig. 4E). In addition, we calculated Moran’s I score, which measured global spatial autocorrelation for genes captured in three sections.

**Fig 4.**
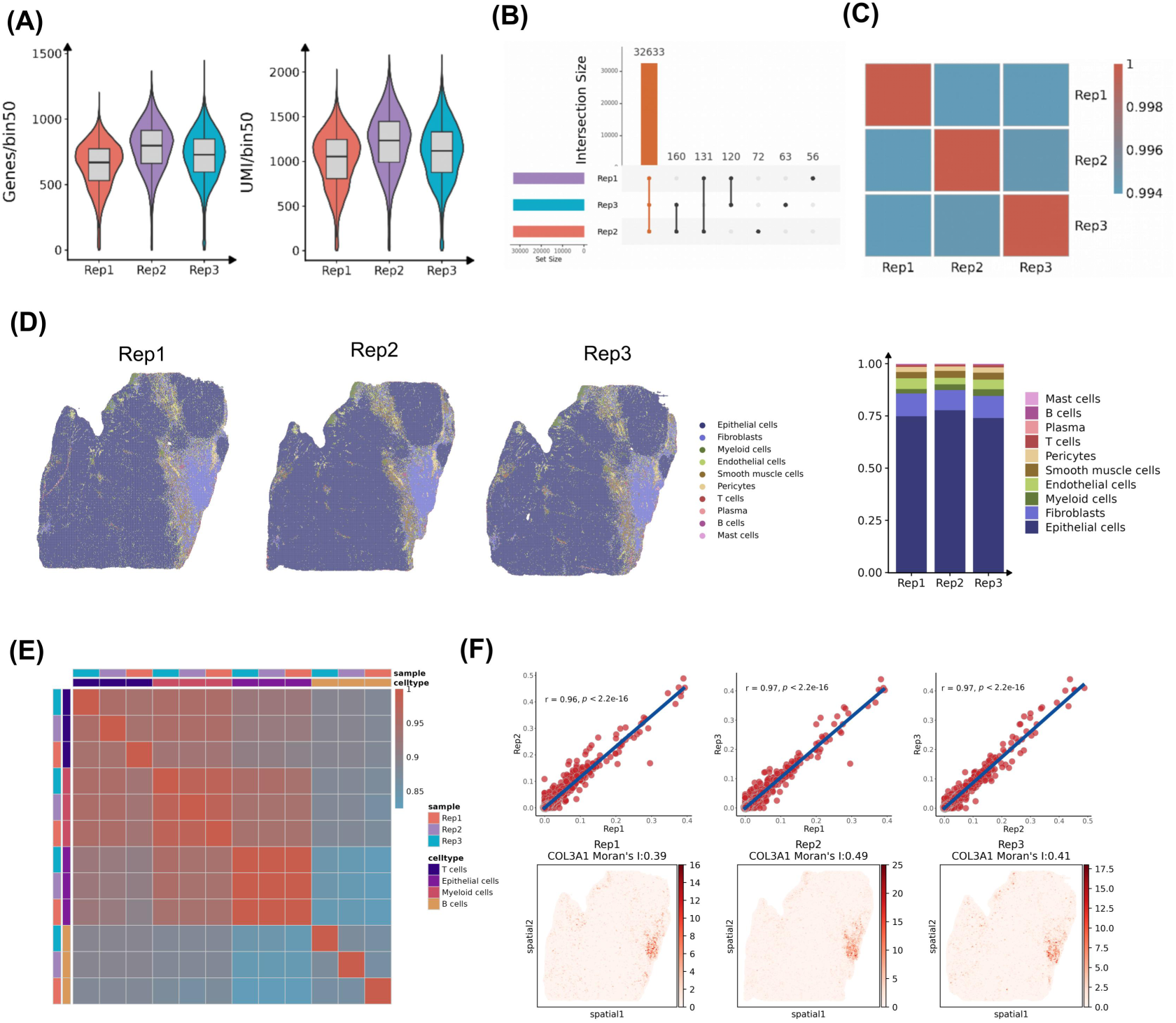
Estimation of reproducibility and stability of ST-FFPE-mIF protocol. (A) Violin plots show the number of captured genes and UMIs at bin50 (50 x 50 DNB) resolution among three consecutive sections. (B) Upset plot shows commonality of captured gene type among three consecutive sections. (C) Correlations of normalized gene expression among three consecutive sections showed stability of the technique. (D) Cell annotation and the proportion of each cell type shows similarities among three consecutive sections. (E) The stability of the technique is demonstrated by the correlations of average gene expression in identical cell types (B, T, Myeloid, Epithelial) across three sections. (F) Scatter plots show a high correlation of the Moran’s I score of the corresponding genes among three sections. The heatmaps show the spatial pattern of *COL3A1* was similar in the three sections.

The correlation analysis of Moran’s I score showed that there were high correlations among the adjacent three sections. Simultaneously, we illustrated the spatial distribution of specific genes such as *COL3A1* across the three sections. Notably, spatial pattern of selected gene expression in all three sections was similar (Fig. 4F and Fig. S2). Altogether, above results suggested that ST-FFPE-mIF had excellent reproducibility.

### ST-FFPE-mIF enables reliable analysis at a single-cell level

We performed cell segmentation on a DAPI-stained image that had been registered with the corresponding ST data. The gene expression matrix at single-cell resolution was generated. Figure 5A showed a representative FOV. The reasonable spatial distribution of annotated cells, specifically expression levels of transcriptional markers in corresponding cell types, and reasonable composition of each cell types indicated that DAPI-based segmentation could achieve accurate cell annotation at single-cell resolution (Fig. 5B-D). Furthermore, to validate results from ST-based cell annotation, we calculated fluorescence signals for each biomarker within individual cells. The results demonstrated that fluorescent signals were correctly expressed in their respective cell types (Fig. 5E-F), further confirming the accuracy of DAPI-staining-based cell segmentation and transcriptome-based cell annotation. Additionally, we inferred spatial intercellular communications between different cell types utilizing StereoSiTE[22] (Fig. S3A). Specific interactions mediated by ligand-receptor genes involved in the EGF pathway primarily occurred between Mast/Myeloid/Endothelial-Epithelial (Fig. S3B). The pattern recognition results exhibited EGF pathway in pattern 3 as an incoming signal of Epithelial (Fig. S3C). Moreover, ST-FFPE-mIF could simultaneously acquire both nuclear and membrane fluorescence signals, thereby making it potential to enhance the precise identification of cell boundaries and accurate cell segmentation. For example, the results of cell segmentation were compared between CD20 staining and DAPI staining in diffuse large B cell lymphoma, which indicated that membrane-based cell segmentation can obtain a relatively reliable cell boundary (Fig. S4).

**Fig 5.**
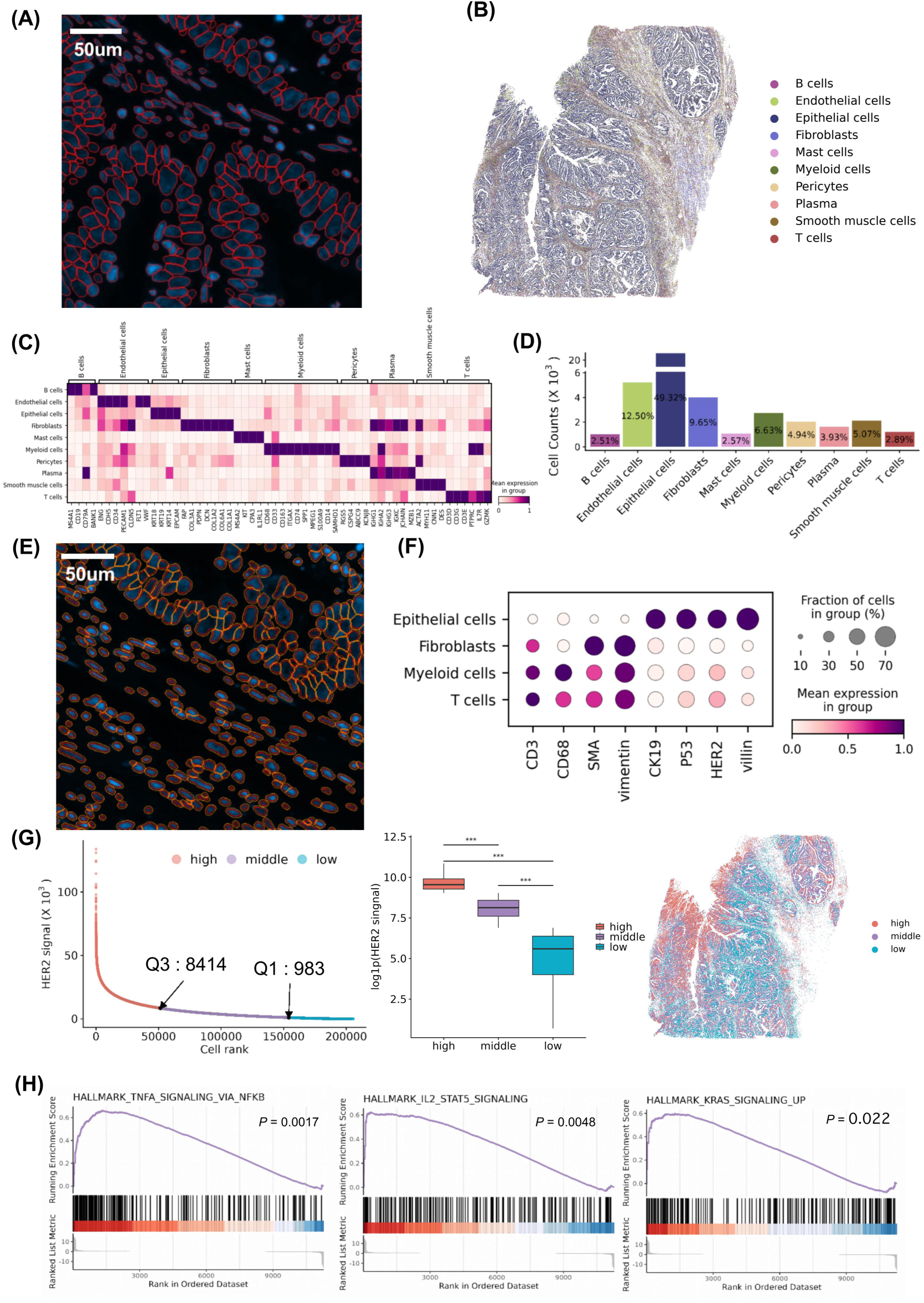
ST-FFPE-mIF enables reliable analyses at single-cell level. (A) Representative FOV illustrates cell segmentation in the DAPI-stained image. The red circles represent the cell boundaries inferred by Cellpose. Scale bar: 50 μm. (B) The spatial visualization displays each cell colored according to its cell type. (C) The heatmap shows specific expression of the classic markers in each cell type. (D) The histogram plot shows the proportion of each cell type. (E) The FOV shows expanded cell boundaries based on the cell segmentation of the DAPI-stained image. The red circles represent cell boundaries inferred by Cellpose. The orange represents expanded boundaries. Scale bar: 50 μm. (F) Dotplot shows that biomarkers were specifically expressed in the corresponding four cell types. (G) Semi-qualitative analysis is performed based on HER2 signal intensity in each cell. The scatter plot shows cell rank based on HER2 signal intensity. The box plot shows there are significant differences among the three groups and the spatial visualization shows spatial distribution of these cell types. (H) GSEA analysis is performed between “high” and “low” groups. Representative figures demonstrate activation of three pathways associated with tumor development.

In addition to validation of cell annotation, mIF images can also be used for semi-qualitative analysis. As a member of the EGFR family of receptor tyrosine kinases, the human epidermal growth factor receptor 2 (HER2) was considered to be a potential therapy target and a prognostic marker in CRC[23]. We calculated HER2 fluorescence signal intensity of all annotated epithelial cells and divided them into three groups according to percentiles, which were named “high” (75th percentile), “middle”, and “low” (25th percentile) (Fig. 5G). Two groups, “high” and “low”, were selected to calculate the difference in gene expression and to perform enrichment analysis on the gene set. GSEA analysis indicated that three pathways “HALLMARK_TNFA_SIGNALING_VIA_NFKB”, “HALLMARK_IL2_STAT5_SIGNALING” and “HALLMARK_KRAS_SIGNALING_UP”, were activated in cells with high HER2 signals (Fig. 5H). These analyses showed that the fluorescence signals of biomarkers can be combined with transcriptomic data to provide clues for exploring tumor development.

## Discussion

Spatial transcriptomics is a start-of-the-art method implemented in many research fields, including cancer biology and developmental biology[24]. Although all proteins are translated from messenger RNAs, post-transcriptional regulation and modifications often lead to discrepancies in their expression patterns[25]. Multi-omics methodologies at spatial resolution are significant for the investigation of the intermolecular/intercellular dynamics on the transcriptome and proteome levels in the same single cells across tumor development and progress. In this paper, we present a co-detection technology that offered subcellular resolution, enabling the simultaneous spatial profiling of the complete transcriptome alongside multiplex immunofluorescence staining.

Herein, we provided a detailed protocol of ST-FFPE-mIF. Taking fluorescent channels, fluorophores, the isotypes, and species reactivity of secondary antibodies into consideration, we proposed an image-based strategy utilizing FL-conjugated primary antibodies together with antibody elution to increase the detection targets and to reduce time-consuming (Fig. 1 and Supplementary 1). Given the accessibility to FL-conjugated primary antibodies, we also tested the traditional secondary antibody-based mIF method and provided the recommended protocol in Supplementary 2. Notably, FL-conjugated primary antibody is frequently regarded as having lower sensitivity for targets with low abundance[26–28]. It is advisable to carefully match fluorescence intensity with antibody signals; for instance, using stronger fluorophores for antibodies that yield weaker signals can effectively enhance the signal-to-noise ratio[29, 30]. It has been reported that multiple rounds of antibody elution have a limited effect on decreased marker intensity or destruction of tissue morphology[26–28]. This indicates that our technology has the potential to increase the target number by raising the number of cycles.

Figure 2-4 proved the authenticity, specificity, sensitivity stability, and reproducibility of ST-FFPE-mIF. To further explore how ST-FFPE-mIF can be used in scientific research, we described the landscape of CRC with ST-FFPE-mIF by inferring spatial intercellular communication at single-cell resolution (Fig. S3A-C). It is important to note that variations in cell morphology and cell size directly lead to the uncertainty of cell boundaries and challenges for accurate segmentation[31–33]. In addition to cell segmentation, the current analytic pipeline also demonstrated two strategies for using mIF images: to identifying the cell types and semi-quantitatively analyzing functional biomarker expression. We found that differential HER2 fluorescence signals were associated with differential gene expression and pathway enrichment. GSEA analysis identified three pathways associated with tumor development[34] that exhibited significant differences in regions characterized by HER2 signal intensities (Fig. 5H).

The current version still has several limitations that need to be addressed in the future. Firstly, we need to improve the iterative cycle number for tagging. It is anticipated that advancements in imaging techniques and an increase in cyclic mIF staining will significantly expand the detection of protein targets. Secondly, it is essential to establish a framework that standardizes the processing and analysis of mIF images. This framework should include multi-round image registration, background noise removal, and quantification of protein expression, utilizing tools such as inForm (Akoya Bioscience, Menlo Park, California, USA) and HALO (Indica Labs, Albuquerque, New Mexico, USA). These considerations will facilitate the creation of a reproducible pipeline for the quantitative assessment of mIF assays in ST-FFPE-mIF.

In summary, the integration of high-precision, unbiased whole transcriptome capture through Stereo-seq-FFPE with multiplex immunofluorescence (mIF) imaging within a single tissue section establishes a robust framework for acquiring spatial multi-omics information at single-cell resolution. This innovative methodology facilitates comprehensive spatial omics analyses, allowing for precise elucidation of structural and functional dynamics within complex multicellular interactions in the tumor microenvironment. The advancements represented by ST-FFPE-mIF highlight its significant potential across diverse applications in fundamental biology, diagnostics, and therapeutic development. Future advancements are expected to provide valuable insights into biological processes, promoting precision medicine and enhancing the ability to address complex health challenges.

## Method

### Sample collection

This study was conducted in strict adherence to ethical standards and guidelines. FFPE samples were sourced from OUTDO BIOTECH CO. LTD (Shanghai, China) through commercial channels. The acquisition and use of human tissue samples strictly adhered to all applicable laws, regulations, and ethical standards. Confidentiality of patient information was upheld at all times. The authors affirm that their search was conducted with integrity and in full compliance with all relevant ethical principles.

### Stereo-seq-mIF solution methodology

#### Tissue de-paraffinization and antigen retrieval

For the human colorectal cancer FFPE sample included in the ST analysis, we performed ST-FFPE-mIF assays using Stereo-FFPE-seq technology (BGI Shenzhen, China) following the manufacturer’s instructions. Tissue sections were cut to a thickness of 5 µm and mounted on Stereo-seq N transcriptomics chips (Cat#211SN114, STOmics). Next, the Stereo-seq N chip with the tissue section was dried and baked overnight at 38°C. Following this, the chip underwent a one-hour baking process at 60°C to facilitate softening and melting of the paraffin. De-paraffinized in Histo-clear substitute and graded ethanol according to Stereo-seq Transcriptomics N Kit (Cat#211SN114, STOmics). Subsequently, the chip with the tissue section was submerged in Tris-EDTA antigen retrieval solution (Cat#C1037, Solarbio) and incubated at 95°C for 25 min to facilitate antigen retrieval.

#### Multiple immunofluorescence staining

After de-paraffinization and antigen retrieval, blocking was performed by adding 100 μL of blocking buffer containing 1x SSC Buffer (Cat#AM9770, Thermo), 0.05 U/μL RNase inhibitor (Cat#M0314L, NEB), 10% 1:1 diluted Gibco™ Horse Serum (Cat#26050070, Thermo) and Gibco™ Goat Serum (Cat#16210064, Thermo), followed by a 20 min incubation at room temperature. After removing the blocking buffer, primary antibodies labeled fluorophores were added at the appropriate concentrations, diluted in the blocking buffer, and incubated for 45 min at room temperature. This was followed by washing the chip with 0.1x SSC Buffer with 0.05 U/μL RNase inhibitor for 2 min, repeated twice. Subsequently, the chip surface was treated with 5 μL of glycerol (Cat#4100854050, Sangon Biotech) to ensure that the tissue was completely covered by glycerol and the coverslip. Imaging was conducted using a ZEISS Axioscan 7 microscope. After imaging, the chip was washed with 0.1 x SSC buffer supplemented with 0.05 U/μL RNase inhibitor. AbEraser (Cat#AXT9710000, AlphaX (Beijing) Biotech Co) was used to elute the antibody from the section. The elution buffer was then applied to the chip surface and incubated at 50°C for 5 min. After removing the solution, the chip was sequentially washed with DMSO (Cat#A610163, Sangon Biotech) and 0.1x SSC Buffer supplemented with 0.05 U/μL RI. For the second round of immunofluorescence staining, the primary antibodies incubation was repeated as previously mentioned. Nuclei were stained with DAPI and performed according to standard procedure. Multispectral images of stained chip were captured using a ZEISS Axioscan 7 microscope. Primary antibodies included alpha -smooth muscle Actin (Cat#S0B1508, Starter, 1:100), Vimentin (Cat#S0B1507, Starter, 1:200), Cytokeratin 19 fragment (Cat#S0B1566, Starter, 1:50), Villin (Cat#S0B1572, Starter, 1:100), p53 (Cat#S0B1569, Starter, 1:200), CD3 epsilon (Cat#S0B0217, Starter, 1:100), CD68 (Cat#S0B1501, Starter, 1:100), HER2 (Cat#S0B1567, Starter, 1:50).

#### Library preparation and sequencing

Sections were processed according to manufacturer’s instructions for the Stereo-seq Transcriptomics N Kit (Cat#211SN114, STOmics). In short, the chip with tissue section was submerged in the FFPE Decrosslinking Reagent for decrosslinking (Cat#211SN114, STOmics). Fixation in pre-cooled methanol (Cat# 34860, Sigma) for 30 min at −20°C ensued. Post-fixation, the Stereo-seq chip was air-dried, and the tissue section was incubated in permeabilization buffer (Cat# 211SN114, STOmics) supplemented with 3 µL of Tetramethylene sulfone (Cat#A610163, Sangon Biotech) for 25 min at 37°C. After permeabilization, the FFPE Dimer mix (Cat#211SN114, STOmics) was added and incubated at room temperature for 40 min. Subsequently, the reverse transcription mix (Cat#211SN114, STOmics) was pipetted onto the chip surface and incubated at 42°C overnight. After reverse transcription, cDNA products were released from the chip by Release Enzyme Mix and collected. Library construction was performed according to the Library preparation Kit (Cat# 111KL114, STOmics) and subsequent DNB generation. Finally, the DNBs were sequenced with the MGI DNBSEQ-T7 or DNBSEQ-G400 Sequencer.

#### H&E stain and immunohistochemistry (IHC)

H&E staining and IHC were performed on adjacent FFPE sections using a standard procedure. Samples were cryo-sectioned at a thickness of 5 µm, and tissue H&E staining was carried out using Hematoxylin (Cat#S330930-2, Dako) and Eosin (Cat#HT110216, Sigma-Aldrich). The staining times varied depending on tissue type. For the biochemical process of IHC, the 4 µm thick section underwent a series of standard procedures including de-paraffinization, hydration, and antigen retrieval. The sections were incubated in 3% hydrogen peroxide to prevent non-specific background staining. The appropriate blocking, antibody incubation, Diaminobenzidine (Cat#DAB-1031, Fuzhou Maixin Biotech) and Hematoxylin (Cat#H8070, Solarbio) stain time were selected according to different sections. The H&E and IHC images were captured with a 20X objective using a ZEISS Axioscan 7 microscope.

#### Stereo-seq-mIF raw data processing

To obtain gene expression matrix from the raw FASTQ data from ST-FFPE-mIF. We aligned the coordination identity sequences (CID) from read 1 to a whitelist of barcodes with known coordinates, allowing for one base mismatch. UMI sequences from read 2 were filtered according to the following criteria: (1) the presence of N bases and (2) having more than two bases with a Phred score lower than 10. Only reads with valid CID and high quality UMI were used for subsequent analyses. The associated read 2 was aligned to the reference genome (GRCh38) using STAR, with only uniquely mapped reads having mapping scores > 10 were kept for gene annotation and read counting. Finally, we counted the deduplicated UMI numbers for each CID and each gene as gene expression counts, and aggregated them to a matrix file with spatial coordinates.

### Staining image registration

#### Two round mIF registration

To register two round mIF images, TrakEM2 within Fiji (v2.16.0)[35] was utilized. The two images were loaded into a new TrarkEM2 project, then one image was locked and the other was adjusted. During the process, we adjusted the position and size so that they matched accurately. Finally, we export two registered flat images.

#### mIF images registration with transcriptome

To register mIF images and transcriptome, Cellbin2 was utilized (https://github.com/STOmics/cellbin2). Initially, the quality of the DAPI-stained image was evaluated. Then, the DAPI-stained image was used for registration with the transcriptome to obtain image transform parameters, which were used to adjust other fluorescence images.

### Quantification and comparison of mIF and IHC stains

To assess the accuracy of signals, we used a pixel-based approach to quantify signals within mIF and IHC stains. The approach was a measure of positive pixel area within a given region. We randomly selected 20 corresponding regions from both mIF and IHC stains using Fiji, distinguished positive and negative pixels based on thresholds, and measured the area. Then, a paired-t test was performed in R software.

### Cell cluster and annotation

We used scanpy (v1.9.3)[36] to construct the data structure and perform cell cluster. The workflow comprised the following operations: (1) Data normalization and scaling; (2) Data dimensionality reduction; (3) Cell clustering using the Leiden algorithm. A single-cell dataset of human CRC[18] was used as a reference to deconvolute a mixture of cells in ST-FFPE-mIF data by Cell2location with N_cell_per_location = 6, detection_alpha = 20. Each cell was assigned a cellular identity. Finally, the proportions of cell types were calculated and visualized in R software.

### Identification of spatially auto-correlated genes

To assess spatial similarity between consecutive sections, we calculated Moran’s I score for captured gene, using scanpy and squidpy (https://github.com/scverse/squidpy). Initially, scanpy was used to filter out cells and genes according to two criteria: (1) Cells with UMI less than 50 and the proportion of mitochondrial genes more than 10%; (2) Genes expressed in lessthan 10 cells. Data normalization and logarithmic transformation were performed after filtering. Finally, squidpy was used to calculate Moran’s I score using the spatial_neighbors and spatial_autocorr functions.

### Cell segmentation

#### DAPI-based cell segmentation

The deep-leaning toolkit Cellpose[32] was used to obtain single-cell spatial coordinates in DAPI-stained images. The built-in “nuclei” model, which is pre-trained to handle segmentation of nuclear fluorescence staining, was used to perform cell segmentation. We set the parameter channel = [3, 0], flow_threshold = 0.8 and diameter = 0.

#### Fluorescence-based cell segmentation

Cellpose was also used to perform cell segmentation in CD20-stained fluorescence images using a built-in “cyto3” model. The parameters were set to channel = [2,0], flow_threshold = 0.8 and diameter = 0.

### Intercellular communications

StereoSiTE (https://github.com/STOmics/StereoSiTE) [22] was downloaded and used to perform spatial cell interaction analysis, The parameters were set to Secreted Signaling = 200, ECM-Receptor = 200, Cell-Cell Contact = 30 The built-in database named “CellChatDB-human.csv” was selected.

### Cell boundary expansion and fluorescence signals calculation

The function “skimage.segmentation.expand_labels” within scikit-image[37] was used to expand the cell boundary. Setting parameter distance = 3. Then, the function “cv2.cvtColor” within opencv (https://github.com/opencv/opencv) was used to transform the RGB image into gray-scale image, and add the gray-scale values of each biomarker within the cell separately to generate matrix.

### GSEA analysis

We used scanpy to extract all epithelial cells and used numpy (https://github.com/numpy/numpy) to calculate the 25th and 75th percentiles of HER2 intensity, which reached 983 and 8414, respectively. The function “scanpy.tl.rank_genes_groups” within scanpy was used to calculate the difference of gene expression. ClusterProfiler (https://github.com/YuLab-SMU/clusterProfiler) was used to perform GSEA analysis, and filter pathways with p-value < 0.05.

## Acknowledge

This study is supported by Science and Technology Innovation Key R&D Program of Chongqing (CSTB2023TIAD-STX0002). The authors would like to acknowledge Miss Tibby Tianbi Duan, Dr. Yu Zhao, Dr. Ying He, Dr. Dan dan Li, and Dr. Mei Li for their kind help. We also wish to thank DCS Cloud (https://cloud.stomics.tech) for providing the computational resources and software support necessary for this study.

## Consent for Publication

The authors declare that the research was conducted in the absence of any commercial or financial relationships that could be construed as a potential conflict of interest.

## Author Contributions

Jiajun Zhang, Xue Zhang, Meng Zhang and Xu Yuan conceived the idea; Jiajun Zhang, Ao Chen, Sha Liao, Xi Feng, Luohao Xu and GuiBo Li supervised the study; Xue Zhang and Meng Zhang designed the experiment; Meng Zhang, Yangyang Song, Xingfeng Yang and Yanan Wu performed the majority of the experiments with the help of Dongsheng Ran, Wenxiao Lei, Hongyan Li and Yongfen Zhang. Yuan Xu. and Xuelin Zhao analyzed the data with the help of Xin Liu, Huaqiang Huang, and Xing Liu provided project support; Xue Zhang, Meng Zhang and Xu Yuan wrote the manuscript; Jiajun Zhang and Xing Liu participated in the manuscript editing and discussion.

---

**Supplementary Fig 1.**
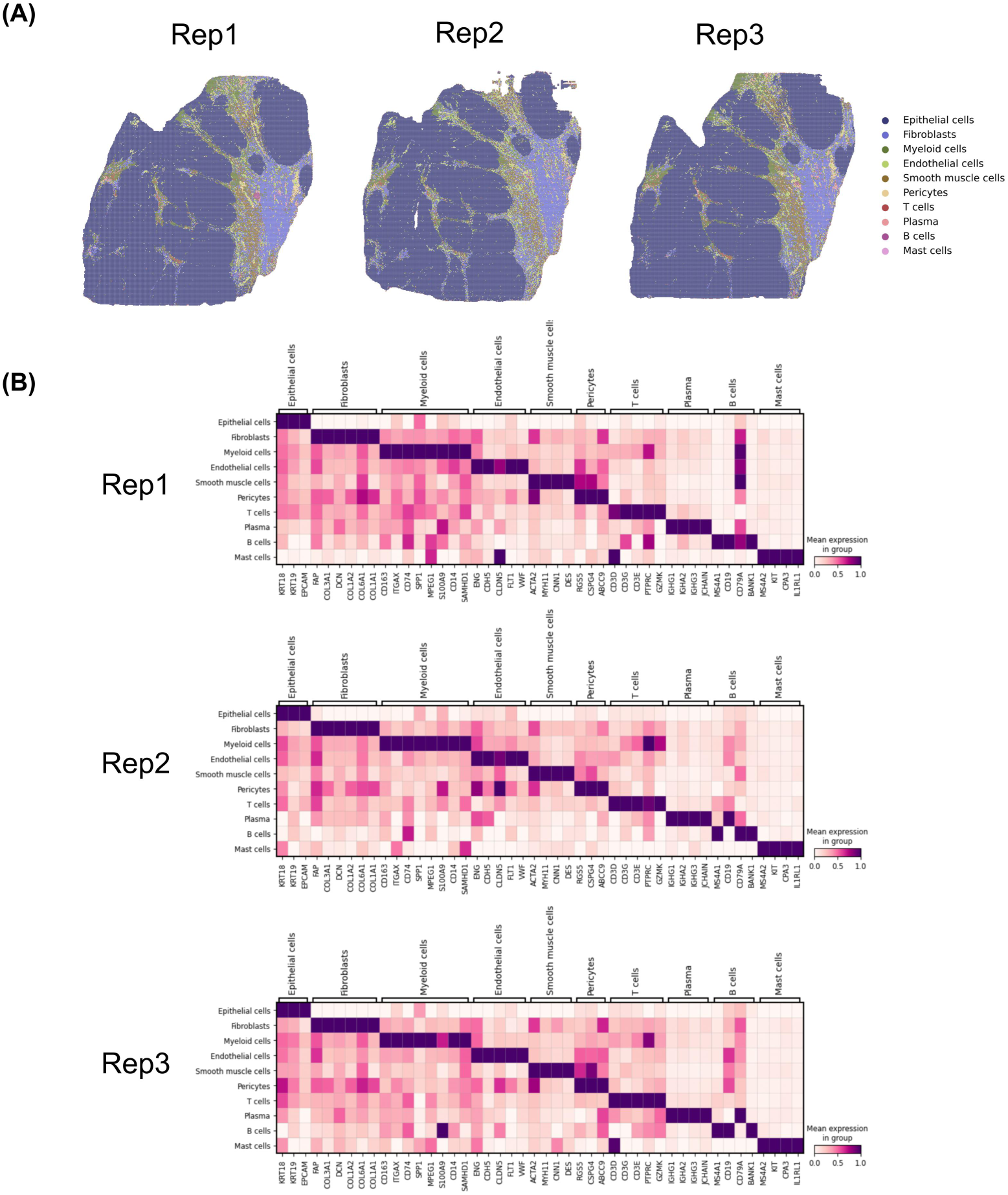
Cell annotation and gene expression of three Stereo-seq-FFPE sections. (A) Spatial maps show the spatial distribution of cell types is similar among three sections generated by the Stereo-seq-FFPE protocol. (B) The heatmaps show scaled expression of classic cell markers is similar among three sections generated by the Stereo-seq-FFPE protocol.

**Supplementary Fig 2.**
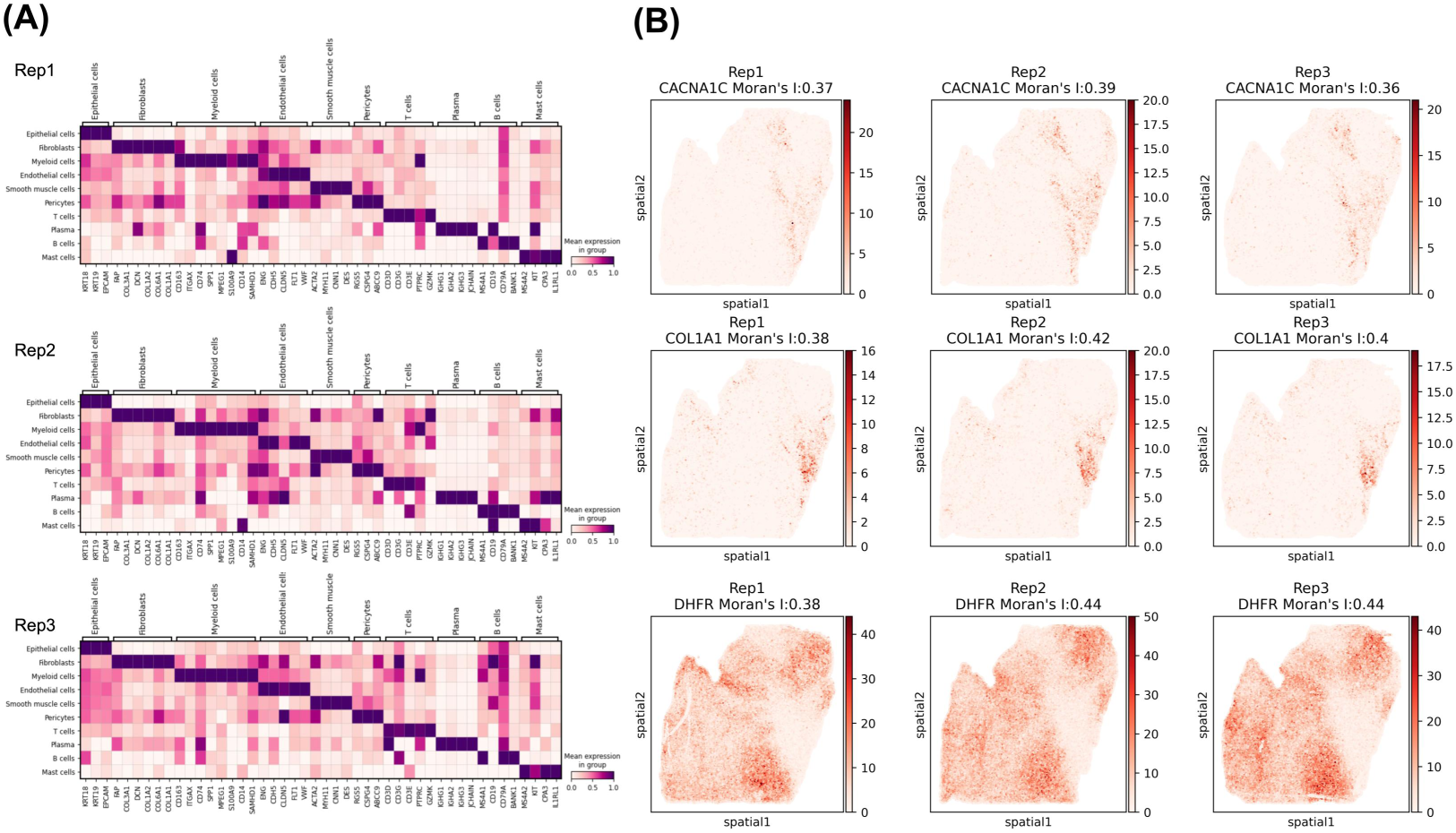
The features of gene expression demonstrated technical reproducibility. (A) The heatmaps illustrate scaled expression of classic cell markers is similar in three replicates. (B) Spatial maps show spatial distribution of genes with high Moran’s I score is similar among the three replicates.

**Supplementary Fig 3.**
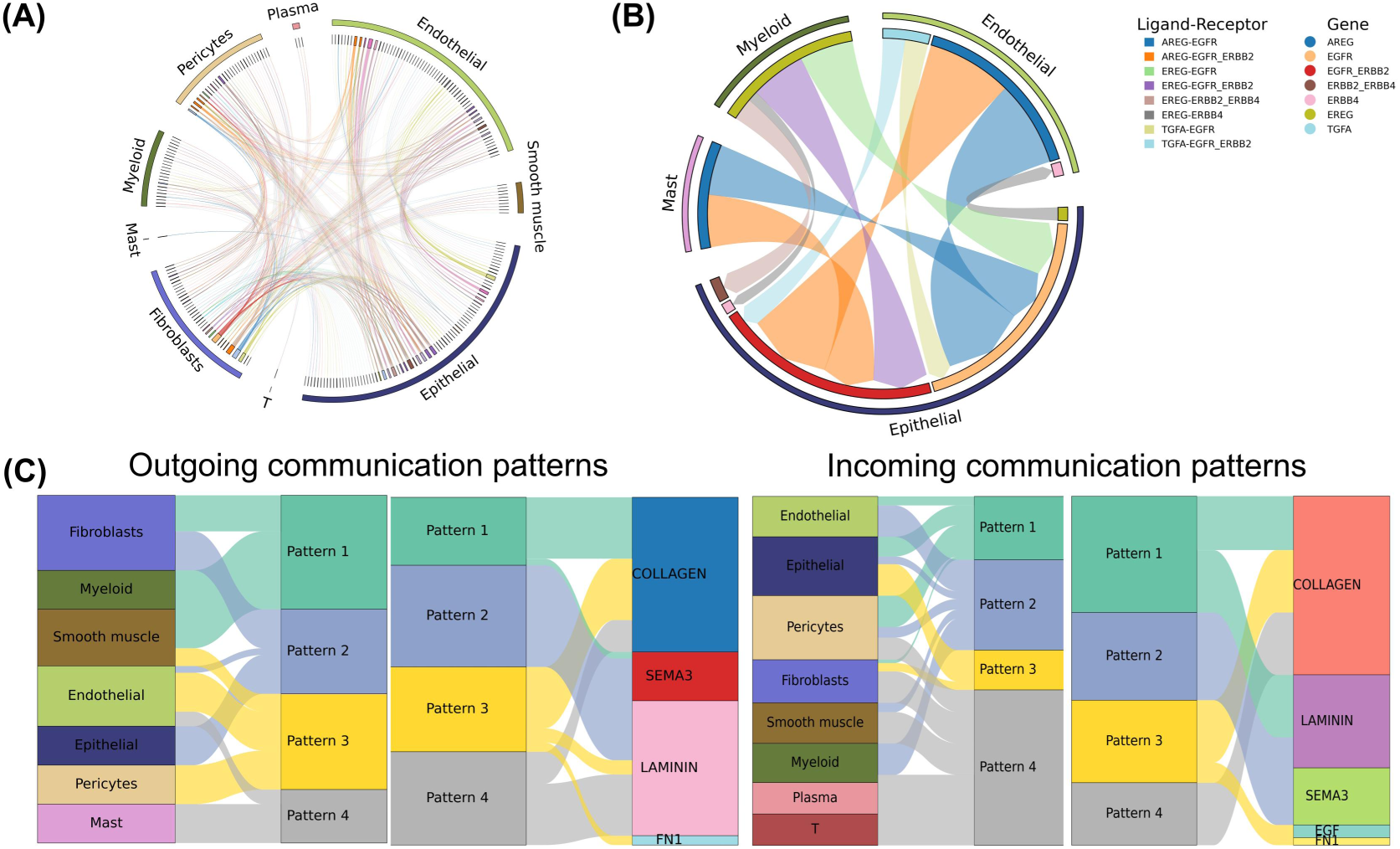
Quality control intercellular communications among annotated clusters in CRC. (A) Circos plot shows cell-to-cell communications activities. (B) Inferred cell-to-cell communication activities associated with EGF signaling pathways explore detailed communications for distinct pathways. (C) The pattern recognition method of cell-to-cell communication pairs demonstrates the coordination of multiple cell groups and signaling pathways in functioning.

**Supplementary Fig 4.**
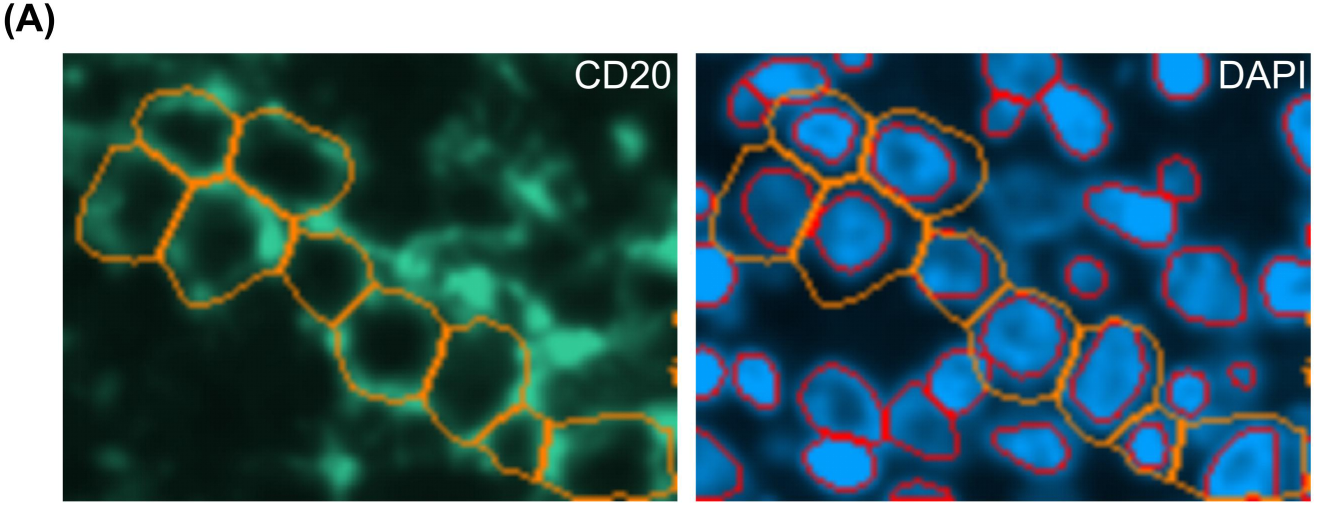
Cell segmentation based on membrane fluorescence signals enables more accurate cell boundary. (A) On the left, the representative FOV represents cell segmentation based on CD20 staining. The orange circles indicate the inferred cell boundaries based on this signal. On the right, red circles indicate the inferred cell boundaries based on DAPI signal.

## Reference

1. Marx, V., Method of the Year: spatially resolved transcriptomics. Nat Methods, 2021. 18(1): p. 9–14.

2. Tian, L., F. Chen, and E.Z. Macosko, The expanding vistas of spatial transcriptomics. Nat Biotechnol, 2023. 41(6): p. 773–782.

3. You, Y., et al., Systematic comparison of sequencing-based spatial transcriptomic methods. Nat Methods, 2024. 21(9): p. 1743–1754.

4. Chen, J., et al., Spatial landscapes of cancers: insights and opportunities. Nat Rev Clin Oncol, 2024. 21(9): p. 660–674.

5. Li, H., et al., A comprehensive benchmarking with practical guidelines for cellular deconvolution of spatial transcriptomics. Nat Commun, 2023. 14(1).

6. Liu, L., et al., Spatiotemporal omics for biology and medicine. Cell, 2024. 187(17): p. 4488–4519.

7. Bressan, D., G. Battistoni, and G.J. Hannon, The dawn of spatial omics. Science, 2023. 381(6657): p. eabq4964.

8. Chen, A., et al., Spatiotemporal transcriptomic atlas of mouse organogenesis using DNA nanoball-patterned arrays. Cell, 2022. 185(10): p. 1777–1792 e21.

9. Chen, K.H., et al., Spatially resolved, highly multiplexed RNA profiling in single cells. Science, 2015. 348(6233).

10. Brady, L., et al., Inter- and intra-tumor heterogeneity of metastatic prostate cancer determined by digital spatial gene expression profiling. Nat Commun, 2021. 12(1).

11. Oliveira, M.F., et al., Characterization of immune cell populations in the tumor microenvironment of colorectal cancer using high definition spatial profiling. bioRxiv, 2024.

12. Liu, Y., et al., High-spatial-resolution multi-omics sequencing via deterministic barcoding in tissue. Cell, 2020. 183(6): p. 1665–1681 e18.

13. Vickovic, S., et al., SM-Omics is an automated platform for high-throughput spatial multi-omics. Nat Commun, 2022. 13(1): p. 795.

14. Harms, P.W., et al., Multiplex immunohistochemistry and immunofluorescence: a practical update for pathologists. Mod Pathol, 2023. 36(7): p. 100197.

15. Schubert, W., et al., Analyzing proteome topology and function by automated multidimensional fluorescence microscopy. Nat Biotechnol, 2006. 24(10): p. 1270–1278.

16. Kleshchevnikov, V., et al., Cell2location maps fine-grained cell types in spatial transcriptomics. Nat Biotechnol, 2022. 40(5): p. 661–671.

17. Mirzazadeh, R., et al., Spatially resolved transcriptomic profiling of degraded and challenging fresh frozen samples. Nat Commun, 2023. 14(1).

18. Pelka, K., et al., Spatially organized multicellular immune hubs in human colorectal cancer. Cell, 2021. 184(18): p. 4734–4752.e20.

19. Rimm, D.L., What brown cannot do for you. Nat Biotechnol, 2006. 24(8): p. 914–6.

20. Monici, M., Cell and tissue autofluorescence research and diagnostic applications. Biotechnol Annu Rev, 2005. p. 227–256.

21. Hoyt, C.C., Multiplex immunofluorescence and multispectral imaging: forming the basis of a clinical test platform for immuno-oncology. Front Mol Biosci, 2021. 8.

22. Liu, X., et al., StereoSiTE: a framework to spatially and quantitatively profile the cellular neighborhood organized iTME. GigaScience, 2024. 13.

23. Mendelsohn, J. and J. Baselga, Epidermal growth factor receptor targeting in cancer. Semin. Oncol, 2006. 33(4): p. 369–385.

24. Williams, C.G., et al., An introduction to spatial transcriptomics for biomedical research. Genome Med, 2022. 14(1).

25. Nerurkar, S.N., et al., Transcriptional spatial profiling of cancer tissues in the era of immunotherapy: the potential and promise. Cancers, 2020. 12(9).

26. Bolognesi, M.M., et al., Multiplex staining by sequential immunostaining and antibody removal on routine tissue sections. J Histochem Cytochem, 2017. 65(8): p. 431–444.

27. Schürch, C.M., et al., Coordinated Cellular Neighborhoods Orchestrate Antitumoral Immunity at the Colorectal Cancer Invasive Front. Cell, 2020. 182(5): p. 1341–1359.e19.

28. Kennedy-Darling, J., et al., Highly multiplexed tissue imaging using repeated oligonucleotide exchange reaction. Eur J Immunol, 2021. 51(5): p. 1262–1277.

29. Lin, J.R., et al., Cyclic Immunofluorescence (CycIF), A highly multiplexed method for single-cell Imaging. Curr protoc chem biol, 2016. 8(4): p. 251–264.

30. Sheng, W., et al., Multiplex immunofluorescence: a powerful tool in cancer immunotherapy. Int J Mol Sci, 2023. 24(4).

31. Israel, U., et al., A foundation model for cell segmentation. bioRxiv, 2024.

32. Stringer, C., et al., Cellpose: a generalist algorithm for cellular segmentation. Nat Methods, 2021. 18(1): p. 100–106.

33. Li, D., et al., Robust blood cell image segmentation method based on neural ordinary differential equations. Comput Math Methods Med, 2021. 2021: p. 5590180.

34. Liberzon, A., et al., The molecular signatures database (MSigDB) hallmark gene set collection. Cell Syst, 2015. 1(6): p. 417–425.

35. Schindelin, J., et al., Fiji: an open-source platform for biological-image analysis. Nat Methods, 2012. 9(7): p. 676–682.

36. Wolf, F.A., P. Angerer, and F.J. Theis, SCANPY: large-scale single-cell gene expression data analysis. Genome Biol, 2018. 19(1).

37. van der Walt, S., et al., scikit-image: image processing in Python. PeerJ, 2014. 2: p. e453.

