## Supplementary 1 for "ST-FFPE-mIF: Integrating Spatial Transcriptomics and Multiplex Immunofluorescence in Formalin-Fixed Paraffin-Embedded Tissues Using Stereo-seq"

### ST-FFPE-mIF User Guide (Direct Fluorescent Antibody Method)

### CHAPTER 1 Introduction

##### 1.1 Product Introduction

The Multiplex Fluorescence (mIF) staining and Formalin-Fixed Paraffin-Embedded (FFPE) Spatial Transcriptomics Co-detection Technique is a cutting-edge methodology that enables the concurrent acquisition of immunofluorescence images for multiple proteins and comprehensive transcriptomic data on a single FFPE tissue section. The mIF process does not interfere with the efficiency of transcriptome capturing. This integration enables a deeper and more comprehensive assessment of valuable biological samples, facilitating the elucidation of intricate pathological or physiological processes. The mIF staining utilizes cyclin-based strategy utilizing FL-conjugated primary antibodies together with antibody elution to increase the detection targets and reduce time consuming, enabling the detection of a protein count commensurate with the channel capacity of a fluorescence microscope within a single staining cycle. Additionally, this technique which does not adversely affect the transcriptomic data, has been tested and demonstrated the capability to achieve the staining efficacy of two rounds through a single elution process.

##### 1.2 Self-Supplied Material List

| | |
|---|---|
| Equipment | Microtome (Leica, Hand-operated rotary slicer)<br>Microtome blade (Leica, Low-profile Blades Disposable, Cat. No. 819) |

| | |
|---|---|
| | Slide warmer/dryer<br>Water Bath (Geyer, Tissue Floating Bath, Lighted)<br>Benchtop centrifuge Microcentrifuge<br>Pipettes<br>PCR (T100 Thermal Cycler, Bio-Rad; ProFle3 x 32-well PCR System, ABI)<br>Constant temperature box<br>NEBNext ® Magnetic Separation Rack for < 200 μL tubes (Cat. No. S1515S)<br>Thermo Fisher Scientific Magnetic rack DynaMagTM-2 for 1.5 ~ 2mL tubes (Cat. No. 12321D)<br>Multi-channel fluorescence microscopes that have stitching functions and both brightfield and fluorescence capacity<br>Benchtop centrifuge<br>Qubit™3 fluorometer (Thermo Fisher, Cat. No. Q33216)<br>Vortex mixer<br>Agilent 2100 Bioanalyzer (Agilent Technologies, Cat. No.G2939AA) |

| | |
|---|---|
| Reagent | Nuclease Free Water (NF water) (Ambion, Cat. No. AM9937)<br>100% Ethanol (Analytical grade)<br>Alexa Fluor series of direct fluorescent antibody<br>DAPI Staining Solution (1 mg/mL) (Thermo Fisher Scientific™ 62248)<br>Horse Serum (Thermo Fisher Scientific Gibco,Cat. No.26050070)<br>Goat Serum (Thermo Fisher Scientific Gibco,Cat. No.16210064)<br>1X TE buffer, pH 9.0 (Ambion, Cat. No. AM9858)<br>Antibody eluent (Absin Bioscience Inc. , Cat. No.abs994 )<br>DMSO (Sangon Biotech, Cat. No. A610163)<br>20X SSC (AMBION, AM9770)<br>VAHTSTM DNA Clean Beads (VAZYME, Cat. No. N411-02)<br>*Agencourt AMPure® XP (Cat. No. A63882)<br>*Beckman Coulter SPRIselect (Cat. No. B23317/B23318/B23)<br>Methanol (SIGMA, Cat. No. 34860-1L-R) |

| | |
|---|---|
| | Hydrochloric acid, HCl (SIGMA, Cat. No. 2104-50ML)<br>Qubit dsDNA HS Assay Kit (Invitrogen, Cat. No. Q32854)<br>High sensitivity DNA kit (Agilent Technologies™, Cat. No. 5067-4626)<br>High sensitivity RNA kit (Agilent Technologies™, Cat. No. 5067-1513)<br>Qubit dsDNA HS Assay Kit (Invitrogen, Cat. No. Q32854)<br>Rnase inhibitor (RI) (Thermo Fisher Scientific , Cat. No. Q33216) |

| | |
|---|---|
| Consumable | Forceps<br>Microscope Slide<br>Blade<br>10 μL、 100 μL、 200 μL、 1,000 μL pipette tips<br>Slide Staining Rack<br>Microscope glass coverslip (area: 18 mm x 18 mm, thickness: 0.13 ~ 0.16 mm)<br>Sterilized Syringe<br>5-Slide Container<br>Microscope Slide Storage Box<br>Microscope Slides or Adhesion Microscope Slides<br>Super PAP Pen (hydrophobic barrier pen)<br>Kimwipes™ delicate task wipes (Kimtech, Cat. No. 34155)<br>Power Dust remover (MATIN, Cat. No. M-6318)<br>1.5 mL、5mL 、15mL、50mL tubes<br>0.2 mL PCR tube (Axygen, Cat. No. PCR-02-C) or (Axygen, Cat. No. PCR-96M2-HS-C)<br>Qubit Assay Tubes (Invitrogen, Cat. No. Q32856) or 0.5 mL tube (Axygen, Cat. No. PCR-05-C)<br>Millex Syringe Filter, Durapore PVDF, 0.22 μm pore size (Millipore, Cat. No. SLGV033N)<br>Corning® 100 mm TC-treated Culture Dish<br>Parafilm (PARAFILM, Cat. No. PM996) |

### Chapter 2 Pre-Experiment: Fluorescent Antibody Validation

The quality of antibodies is pivotal for the success of the experiment. We recommend testing antibody efficacy on positive tissue controls to ensure proper fluorescent labeling and clear signal detection. Additionally, it's important to check the overall staining pattern by placing antibodies with different fluorophores on the same slide before the formal experiment. Finally, confirming the specificity of the fluorescent antibodies by comparing their images with those of the corresponding chromogenic immunohistochemistry is essential.

#### 2.1 Experiment Preparation

Unless otherwise specified, use nuclease-free water (NF-$H_2O$) to prepare all reagents being.

Prepared prior to this experiment.

| Reagent/equipment | Section Preparation Steps | Maintenance |
|---|---|---|
| 400mL 30% ethanol | Dilute anhydrous ethanol to 30% using dd$H_2O$ | Room temperature up to 1 day |
| 100mL 96% ethanol | Dilute anhydrous ethanol to 96% using dd$H_2O$ | Room temperature up to 1 day |
| 50mL 90% ethanol | Dilute anhydrous ethanol to 90% using dd$H_2O$ | Room temperature up to 1 day |
| 50mL 80% ethanol | Dilute anhydrous ethanol to 80% using dd$H_2O$ | Room temperature up to 1 day |
| 50mL 70% ethanol | Dilute anhydrous ethanol to 70% using dd$H_2O$ | Room temperature up to 1 day |
| 50mL 50% ethanol | Dilute anhydrous ethanol to 50% using dd$H_2O$ | Room temperature up to 1 day |
| 50 mL 30% ethanol | Dilute anhydrous ethanol to 30% using dd$H_2O$ | Room temperature up to 1 day |
| 1X SSC | Add 1mL 20X SSC into 20 mL dd$H_2O$ and mix well. | Room temperature up to 1 day |

| | | |
|---|---|---|
| Tris-EDTA (PH 9.0) Antigen Retrieval Buffer | For each 5-slide container, prepare at least 20 mL solution and place it in the container before use. Preheat the solution in the 95°C water bath for 5 min | 4°C |
| Microtome | 5 μm is recommended for normal tissues and 4 μm for tissues with high fat content (such as breast cancer ) | - |
| Slide warmer/dryer | 42°C / 60°C | - |
| Water Bath | 42°C | - |
| Heating circulating water bath | 95°C | - |
| Shaker | 200 ~ 300 r/min | - |
| Fluorescence microscopy | stitching functions and both bright field and fluorescence capacity | - |

##### 2.2 Tissue Sectioning and Mounting

***※ This procedure should be carried out by technicians who are experienced in performing paraffin sectioning***

a. Make sure your water bath and baking machine have been turned on and set to 39 ~ 42°C and 42°C, respectively (If a different water bath temperature has been employed at your institution's pathology/histology laboratories, always adhere to the established protocols);

b. Prepare the microtome, histology brushes, forceps, new microtome blades, and a container filled with 300 ~ 400 mL of 30% ethanol;

c. Place the paraffin block face down in an ice-bath for 10 ~ 30 min, or cool the tissue surface on a cooling platform for 5 ~ 10 min;

d. Trim the block to expose enough tissue surface from which a representative section can be cut. Trimming is normally done at a thickness of 10 ~ 30 μm.
e. Adjust the section thickness to 5 μm for regular tissue and 4 μm for high–fat–content tissue to reduce the possibility of section detachment in subsequent operations.
f. The number of sections is related to the number of antibodies to be detected. If detecting 8 fluorescently labeled antibodies, it is necessary to cut 9 to 10 consecutive sections. Mark the order of the sections in pencil on the blank area of the slide. Of these, the 4 sections at the beginning and end are used for immunohistochemical staining with naked antibodies corresponding to the fluorescent antibodies, while the middle 1 to 2 sections are used for multiplex immunofluorescence staining.

***※: A paraffin section is divided into two sides, side A and side B. The side facing the microtome is side A (matte side), and the side touching the blade is side B (smooth side). Always keep side B facing down when it is floating in the water bath, or keep it mounted onto the microscopic glass slide to prevent the section detachment.***

g. Carefully transfer the section with side B facing down to the container filled with 30% ethanol using a histology brush or clean forceps.

**This step is to assist in transferring the sections to the water bath and ensures smooth section spreading. If performed by an experienced technician, this step can be skipped.**

h. Use a clean microscopic glass slide to pick up the section and float it on the surface of the water in the preheated water bath. Make sure side B is facing down.
i. After the tissue sections are completely flattened, use Adhesion Microscope Slides to adhere the tissue to the center of the slide, ensuring that side B is in

contact with the slide (it is not recommended to attach two or more tissues on a single slide). After the sections are picked up, gently shake off the excess water around the tissue.

Notes:

- Technicians with extensive sectioning experience may follow their own practices for sectioning and mounting operations.
- After mounting, care should be taken to ensure that the surface of the sections adhered to the slides is free of air bubbles.
- Slide warming can also be done using a metal bath or any other module capable of providing accurate and consistent heating.

##### 2.3 De-paraffinization

a. Set aside the ethanol of different concentrations and $ddH_2O$ you prepared in 2.1 Experiment Preparation, turn on the Slide Dryer in advance (or Water Bath-Slide Dryer, metal bath, or PCR thermal cycler with PCR Adaptor), and set the baking temperature to 60˚C.

b. Transfer the tissue-mounted slides onto the Slide Dryer and bake at 60˚C for 1 hr.

c. Prepare the following reagents in staining jars, at least 30 mL for each jar:

| Description Quantity | Quantity |
|---|---|
| Histo-clear | 2 |
| 100% ethanol | 2 |
| 90% ethanol | 2 |
| 70% ethanol | 1 |
| 50% ethanol | 1 |
| 30% ethanol | 1 |
| $ddH_2O$ | 1 |

##### Xylene can be used in place of Histo-clear

d. After baking, place the slides into Histo-clear at room temperature for 20 min, then transfer it to Histo-clear at room temperature for another 20 min.

e. Take the slides out from Histo-clear and remove the excess Histo-clear solution with dust-free paper.

f. Sequentially place the slides into the following containers:

   1. 100% ethanol for 5 min
   2. 100% ethanol for 5 min
   3. 96% ethanol for 5 min
   4. 96% ethanol for 5 min
   5. 90% ethanol for 2 min
   6. 80% ethanol for 2 min
   7. 70% ethanol for 2 min
   8. 50% ethanol for 2 min
   9. 30% ethanol for 2 min
   10. dd $H_2O$ for 1 min.

g. Take the slides out and remove excess dd$H_2O$ from around and the back of the slide with dust-free paper without touching the tissue.

#### 2.4 Antigen retrieval

a. Prepare a 95°C water bath in advance. Place Tris-EDTA (pH 9.0) antigen retrieval solution in a 5-slide container and preheat it in the 95 ° C water bath for 5 min;

b. Remove the preheated antigen retrieval solution from the water bath. Gently place the slides into a 5-slide container using tweezers, and then immerse the container in the 95°C water bath for 20 min.

c. During the antigen retrieval waiting period, refer to Table 2-1 to prepare the reagents needed for the subsequent blocking and antibody incubation steps.

d. Remove the 5-slide container from the water bath and allow it to cool down gradually at room temperature.

Table 2-1 Reagent preparation for tissue blocking and antibody incubation

| Reagent | Preparation Steps | Maintenance |
|---|---|---|
| Mixed serum | Retrieve aliquoted goat serum and horse serum from -20°C storage. Allow them to thaw and then mix them in 1:1 ratio in a tube. | On ice until use |
| Direct fluorescent antibody | Take them out of -20°C or 4°C (depending on the manufacturer's instruction) and centrifuge at 14,000g, 4°C for 10 min then leave them on ice. | On ice until use |
| Diluted antibody (optional) | Primary antibodies can be diluted with blocking solution to a desired concentration if needed. | On ice until use |
| DAPI | Take them out of -20°C or 4°C (depending on the manufacturer's instruction) and leave them on ice. | On ice until use |

Notes:

- Tris-EDTA is a commonly used antigen retrieval method, but antigen retrieval can also be performed according to the specific requirements of each antibody.
- Antigen retrieval solution can be placed in containers other than the 5-slide container, depending on the situation.
- If using 5-slide containers as containers for antigen retrieval solution, the number of containers needed should be determined based on the number of slides.

e. Remove the first and last four sections and proceed with the routine immunohistochemistry (IHC) protocol.

#### 2.5 Tissue Blocking and Antibody Incubation

a. After the antigen retrieval solution has cooled down, use forceps to remove the slides and place them into a staining jar that has been pre-filled with 1X SSC. Wash the slides on a shaker at room temperature incubate for 3 min and then discard. Repeat this step three times.

b. Take out the microscope slide and wipe off excess liquid from around and the back of the slide with dust-free paper without touching the tissue.

c. Use a Super PAP Pen (hydrophobic barrier pen) on the microscope slide to draw a circle around the tissue to create a hydrophobic exclusion zone to prevent the subsequent addition of fluid from escaping.

d. Prepare the blocking solution according to Table 2-2 below. The amount of blocking solution to prepare is 200 μL per slide, with 100 μL used for blocking and the other 100 μL used for preparing the antibody incubation solution. Add 100 μL/slide of blocking solution to hydrophobic exclusion zone surrounding the tissue. Adjust the volume according to the size of the hydrophobic circle (for example, if the circle is 0.5 cm X 0.5 cm, use 50 μL per slide), and incubate at room temperature for 20 minutes.

Table 2-2 blocking solution (one slide)

| Reagent | 1X (μL) |
|---|---|
| 1X SSC | 180 |
| Mixed serum | 20 |
| Total | 200 |

e. While waiting for the incubation to be done, prepare the antibody solution according to Table 2-3. For the initial determination of antibody dosage, prepare the antibody solution according to the concentration recommended by the manufacturer. If the recommended dilution is 1:200, for example, add 0.5 μL of antibody. Mix the antibody incubation solution by pipetting, and then place it on ice for use.

Table 2-3 antibody solution (one slide)

| Reagent | 1X (μL) |
|---|---|
| Direct fluorescent antibody 1 | V1 |
| Direct fluorescent antibody 2<br>……<br>Direct fluorescent antibody n | V2<br>……<br>Vn |
| Blocking solution | 100 – (V1+V2+…+Vn) |
| Total | 100 |

f. Discard the blocking solution with a pipette. For experimental groups: Slowly add the antibody solution from the non-tissue area until the solution covers the tissue section. Do not exceed 100 μL/slide. Make sure to label the dilution ratio on the microscope slide. Incubate at room temperature for 45 min in the dark. For the negative control group: Add 100 μL/slide of blocking solution. Incubate at room temperature for 45 min in the dark.

※ ***Make sure not to dry the tissue during the solution addition process. Otherwise, dried tissue might generate non-specific signals which interferes with the final results.***

#### 2.6 DAPI Staining

a. While waiting for the antibody incubation to be done, prepare DAPI staining solution according to Table 2-4. Mix with pipette then leave it on ice in the dark until use;

b. Discard the antibody solution with a pipette;

c. Wash by adding 100 μL/slide of 1X SSC. Incubate for 3 min and then discard. Repeat c. twice for a total three-time wash;

d. Slowly add 100 μL/slide of DAPI staining solution to the tissue. Incubate for 2 min at room temperature in the dark;

e. Discard the DAPI staining solution with a pipette. Wash by adding 100 μL/ slide of 1X SSC then discard;

f. Air dry the microscope glass slides or dry with a hand-held fan until no residual solution is left on the tissue;

g. Pipette 5 μL glycerol/slide gently onto the center of the tissue without introducing air bubbles. With a pair of forceps, place one end of the coverslip onto the glycerol covered tissue while holding the other end and then gradually lower the coverslip. Proceed to image immediately.

Table 2-4 DAPI staining solution (one slide)

| Reagent | 1X (μL) |
|---|---|
| 5X SSC | 60 |
| 50X-diluted DAPI solution | 2 |
| Nuclease-free water | 38 |
| Total | 100 |

#### 2.7 Imaging

a. Take fluorescent images with microscopes that have stitching functions and both bright field and fluorescence capacity.

b. Required fluorescence channels will depend on antibody selection. Exposure times will depend on antibody sensitivity and fluorophores. Adjust the focus, light intensity, and exposure time based on image clarity. Scan using 10X or 20X lens.

※ ***The photography parameters for the negative control must be consistent with those of the experimental group.***

#### Guidelines for Selecting Optimal Antibody Concentration

The principle of optimal antibody concentration selection is to select the antibody concentration that results in the best fluorescent signal of desired cells while minimizing nonspecific background staining.

#### mIF Pilot Experiment Results

A successful mIF pilot experiment should ensure that each channel can obtain the specific staining result as expected, the signal-to-noise ratio is maintained at a reasonable intensity, and there is no significant fluorescence bleed-through among different channels which can affect the subsequent analysis.

#### Staining Accuracy Assessment

If the pattern of the positive signal expression of the fluorescent antibody matches that of the "gold standard" immunohistochemistry (IHC) and the positive signal rate is highly consistent (due to the use of adjacent sections, the signal consistency rate cannot be completely identical), then the fluorescent antibody can be considered to specifically bind to the corresponding target antigen, indicating high staining accuracy.

### CHAPTER 3 Sample Preparation

For paraffin sample sectioning and mounting guides, refer to **Sample Preparation, Sectioning, and Mounting Guide for Formalin-fixed and Paraffin-embedded (FFPE) Samples on Stereo-seq Chip Slides (Document No.: STUM-SP003).**

Note：This guide explained how to check the RNA quality (DV200 value) of a FFPE tissue sample before proceeding to the Stereo-seq experiment. It is strongly recommended that you proceed only with tissue samples with a DV200 ⩾ 30%.

### CHAPTER 4 mIF-Stereo-seq solution for FFPE operating procedure

#### 4.1 Experiment Preparation (one slide)

Unless otherwise specified, use nuclease-free water to prepare all reagents being prepared prior to this experiment.

| Prep Day | Reagent | Section Preparation Steps | Maintenance |
|---|---|---|---|
| Day 1 | 400 mL 30% ethanol | Dilute anhydrous ethanol to 30% using $ddH_2O$ | Room temperature up to 1 day |
| Day 2 | 100mL 96% ethanol | Dilute anhydrous ethanol to 96% using $ddH_2O$ | Room temperature up to 1 day |
| | 50 mL 90% ethanol | Dilute anhydrous ethanol to 90% using $ddH_2O$ | Room temperature up to 1 day |
| | 50 mL 80% ethanol | Dilute anhydrous ethanol to 80% using $ddH_2O$ | Room temperature up to 1 day |
| | 50 mL 70% ethanol | Dilute anhydrous ethanol to 70% using $ddH_2O$ | Room temperature up to 1 day |
| | 50 mL 50% ethanol | Dilute anhydrous ethanol to 50% using $ddH_2O$ | Room temperature up to 1 day |
| | 50 mL 30% ethanol | Dilute anhydrous ethanol to 30% using $ddH_2O$ | Room temperature up to 1 day |
| | Tris-EDTA (PH 9.0) Antigen Retrieval Buffer | Prepare at least 400 μL /chip of solution and place it in an EP tube 5 minutes before use. Preheat the solution in a 90°C PCR heating block. | 4°C |
| | Aliquot Mix Serum | Retrieve aliquoted goat serum and horse serum from -20°C storage. Thaw serum then filter it with a 0.22 μm. Allow goat serum and horse serum to thaw | On ice until |

| | | | |
|---|---|---|---|
| | | and then mix them in 1:1 ratio in a tube | |
| | Direct fluorescent antibody | Take them out of −20°C or 4°C (depending on the manufacturer's instruction) and centrifuge at 14,000g, 4°C for 10 min then leave them on ice. | On ice until use |
| | DAPI | Take them out of −20°C or 4°C (depending on the manufacturer's instruction) and leave them on ice. | On ice until use |
| | 5X SSC | Add 12.5 mL 20X SSC into 37.5 mL dd$H_2O$ and mix well | Room temperature up to 1 day |
| | 1X SSC (with 5% RI) | 800 μL/ slide (760 μL 0.1X SSC + 40 μL RI) | On ice until use |
| | 0.1X SSC | Add 100 μL 20X SSC into 19.9 mL dd$H_2O$ and mix well | Room temperature up to 1 day |
| | 0.1X SSC (with 5%RI) | Add 0.5 mL RI into 9.5 mL 0.1X SSC | Room temperature up to 1 day |
| | Antibody elution Buffer | Prepare at least 400 μL /chip of solution and place it in an EP tube 5 minutes before use. Preheat the solution in a 50°C PCR heating block. | 4℃ |
| | FFPE Decrosslinking Reagent | Take it out of −20°C in advance, equilibrate to room temperature until it thaws, then mix well to ensure that no white precipitates are visible. | Room temperature |
| | If white precipitates are visible in the reagent, dissolve them by heating | | |

| | | |
|---|---|---|
| the buffer at > 50°C and equilibrate to room temperature before mixing. | | |
| 0.01 N HCl Permeabilization | Prepare at least 1 mL of 0.01 N HCl per sample. Configure HCl to 0.01 N. Measure and make sure the pH = 2. | Room temperature for 48 hr (Storing longer than 48 hr will affect the desired pH. Please use within 48 hr of preparation) |
| **Always use freshly prepared 0.01 N HCl (pH = 2.0±0.1). For pre-made 0.1 N HCl and newly purchased HCl, check the pH prior to the experiments.** | | |
| 10X Permeabilization Reagent Stock Solution | Add 1 mL freshly prepared 0.01 N HCl to dissolve PR Enzyme, and thoroughly mix the reagent through pipetting. | -20°C up to 1 month |
| **DO NOT vortex the permeabilization enzyme. Mix by pipette before using.**<br>**Aliquot this 10X stock solution to avoid freeze-thaw cycles.** | | |
| 1X Permeabilization Reagent Solution | Make 1X PR solution (200 μL / chip) by diluting 10X PR stock solution with 0.01N HCl. | On ice until use, up to 6 hr |
| FFPE RT Buffer Mix | Take it out of -20°C in advance, equilibrate to room temperature until it thaws, then mix well to ensure that no white precipitates are visible. | On ice until use |
| FFPE Dimer | Take it out of -20°C in advance, and thaw on ice. | On ice until use |
| FFPE RT Oligo | Take it out of -20°C in advance, and thaw on ice. | On ice until use |
| cDNA Release Buffer | Take it out in advance and heat the buffer for 5 min | |

| | | | |
|---|---|---|---|
| | | at 55°C to dissolve the precipitate. Equilibrate it to room temperature prior to use. | Room temperature |
| **If white precipitates are visible in the buffer, dissolve them by heating the buffer at 55° C again and equilibrate to room temperature before mixing.** | | | |
| Day 3 | Magnetic Beads cDNA Purification | Take it out in advance and equilibrate to room temperature at least 30 min prior to use. | 4°C |
| | 10 mL 80% Ethanol | Dilute 100% ethanol to 80%. | Room temperature up to 1 day |
| | cDNA Amplification Mix | Take it out of −20°C in advance, and thaw on ice. | On ice until use |
| | FFPE cDNA Primer Mix | Take it out of −20°C in advance, and thaw on ice. | On ice until use |
| | TE buffer, pH 8.0 | Set it aside at room temperature until use | Room temperature |

**Other Preparation**

| Equipment | Set up | Notes |
|---|---|---|
| PCR Thermal Cycler | Set the temperature in the following order:<br>1. 90°C for antigen retrieval (heated lid at 85°C ).<br>2. 95°C for decrosslinking (heated lid at 85°C).<br>3. 37°C for slide baking, permeabilization, and RT ligation (heated lid at 42°C).<br>4. 42°C for slide baking and RT + ligation (heated lid at 42°C).<br>5. 55°C for cDNA release (heated lid at 60°C) | Check the PCR thermal cycler for Any abnormalities, and replace it if necessary. |
| Fluorescence Microscope | Microscopes that have stitching functions and both bright field and fluorescence capacity. | Please select the channels according to the fluorescent emission wavelengths of **Your Antibodies**. |

#### 4.2 Tissue Sectioning and Mounting

***※ This procedure should be carried out by technicians who are experienced in performing paraffin sectioning***

a. Make sure your water bath and baking machine have been turned on and set to 39 ~ 42°C and 38°C, respectively (If a different water bath temperature has been employed at your institution's pathology/histology laboratories, always adhere to the established protocols);

b. Prepare the microtome, histology brushes, forceps, new microtome blades, and a container filled with 300 ~ 400 mL of 30% ethanol;

c. Place the paraffin block face down in an ice-bath for 10 ~ 30 min, or cool the tissue surface on a cooling platform for 5 ~ 10 min;

d. Trim the block to expose enough tissue surface from which a representative section can be cut. Trimming is normally done at a thickness of 10 ~ 30 μm

e. Adjust the section thickness to 5 μm for regular tissue and 4 μm for high-fat-content tissue to reduce the possibility of section detachment in subsequent operations.

***※ A paraffin section is divided into two sides, side A and side B. The side facing the microtome is side A (matte side), and the side touching the blade is side B (smooth side). Always keep side B facing down when it is floating in the water bath, or keep it mounted onto the microscopic glass slide to prevent the section detachment.***

f. Carefully transfer the section with side B facing down to the container filled with 30% ethanol using a histology brush or clean forceps.

> **This step is to assist in transferring the sections to the water bath and ensures smooth section spreading. If performed by an experienced technician, this step can be skipped.**

g. Use a clean microscopic glass slide to pick up the section and float it on the surface of the water in the preheated water bath. Make sure side B is facing down.

h. After the tissue sections are completely flattened, use Stereo-seq Chip Slide to adhere the tissue to the center of the chip, ensuring that side B is in contact with the chip (it is not recommended to attach two or more tissues on a single chip). After the sections are picked up, gently shake off the excess water around the tissue.

i. Transfer the tissue-mounted Stereo-seq Chip Slide onto the Slide Dryer and bake it at 37 ~ 40℃ overnight. Do not exceed 16 hr.

Notes:

- Technicians with extensive sectioning experience may follow their own practices for sectioning and mounting operations.
- After mounting, care should be taken to ensure that the surface of the sections adhered to the chip is free of air bubbles.
- Slide warming can also be done using a metal bath or any other module capable of providing accurate and consistent heating

##### 4.3. De-paraffinization

a. Set aside the ethanol of different concentrations and dd$H_2O$ you prepared in 4.1 Experiment Preparation, turn on the Slide Dryer in advance (or Water bath-Slide Dryer, metal bath, or PCR thermal cycler with PCR Adaptor), and set the baking temperature to 60˚C.

b. Transfer the tissue-mounted Stereo-seq Chip Slide onto the Slide Dryer and bake at 60˚C for 1 hr.

**c.** Prepare the following reagents in staining jars, at least 30 mL for each jar:

| Description Quantity | Quantity |
|---|---|
| Histo-clear | 2 |
| 100% ethanol | 2 |
| 90% ethanol | 2 |
| 70% ethanol | 1 |
| 50% ethanol | 1 |
| 30% ethanol | 1 |
| dd $H_2O$ | 1 |

**Xylene can be used in place of Histo-clear**

d. After baking, place the Stereo-seq Chip Slide into Histo-clear at room temperature for 20 min, then transfer it to Histo-clear at room temperature for another 20 min.

e. Take the Stereo-seq Chip Slide out from Histo-clear and remove the excess Histo-clear solution with dust-free paper.

f. Sequentially place the Stereo-seq Chip Slide into the following containers:
   1. 100% ethanol for 5 min
   2. 100% ethanol for 5 min
   3. 96% ethanol for 5 min
   4. 96% ethanol for 5 min
   5. 90% ethanol for 2 min
   6. 80% ethanol for 2 min
   7. 70% ethanol for 2 min
   8. 50% ethanol for 2 min
   9. 30% ethanol for 2 min
   10. dd $H_2O$ for 1 min.

g. Take the Stereo-seq Chip Slide out and remove excess dd$H_2O$ from around and the back of the slide with dust-free paper without touching the chip.

#### 4.4 Antigen Retrieval

a. Place the PCR Adaptor in the PCR thermal cycler and set the program as follows:

Table 4-1 Program selection: Antigen retrieval

| Temperature | Time |
|---|---|
| (Heated lid) 85°C | on |
| 90°C | ∞ |

b. Preheat the Tris-EDTA (pH 9.0) antigen retrieval solution on the PCR instrument 20 minutes in advance, 400 μL /chip;

c. Assemble the Cassette and Gasket then place the Stereo-seq Chip Slide in the Cassette according to the guide written in 'Stereo-seq TRANSCRIPTOMICS SET FOR FFPE User Manual Appendix I': Stereo-seq Slide Cassette Assembly. It is recommended that you practice with a regular blank glass slide. Grip along the Stereo-seq Cassette to make sure the Stereo-seq Chip Slide has been locked in place;

d. Add 400 μL Tris-EDTA (PH 9.0) into the well of the Stereo-seq Slide Cassette. Apply sealing tape to the Stereo-seq Slide Cassette and make sure it is sealed tightly;

e. Place the Stereo-seq Slide Cassette in the 90°C PCR thermal cycler and let the chip incubate inside the PCR thermal cycler at 90°C for 25 min.;

f. During the antigen retrieval waiting period, refer to Table 4-2 to prepare the reagents needed for the subsequent blocking and antibody incubation steps;

Table 4-2 Reagent preparation for tissue blocking and antibody incubation

| Reagent | Preparation Steps | Maintenance |
|---|---|---|
| Mixed serum | Retrieve aliquoted goat serum and horse serum from -20°C storage. Allow them to thaw and then mix them in 1:1 ratio in a tube | On ice until use |
| Direct fluorescent antibody | Take them out of -20°C or 4°C (depending on the manufacturer's instruction) and centrifuge at 14,000g, 4°C for 10 min then leave them on ice. | On ice until use |

| | | |
|---|---|---|
| Diluted antibody (optional) | Primary antibodies can be diluted with blocking solution to a desired concentration if needed. | On ice until use |
| DAPI | Take them out of -20°C or 4°C (depending on the manufacturer's instruction) and leave them on ice. | On ice until use |

#### 4.5 Blocking and Antibody Incubation

a. When incubation is completed, remove the Stereo-seq Slide Cassette from the PCR Adaptor (90°C) and allow it to cool down gradually at room temperature;

b. Carefully transfer the Stereo-seq Slide Cassette to the nearest bench and peel off the sealing tape. Discard the antigen retrieval buffer with a pipette;

c. Remove the slide from the Stereo-seq Slide Cassette according to instructions in Appendix I: Stereo-seq Slide Cassette Assembly, discard the cassette and gasket, then immediately proceed to the next step;

※ ***The cassette and gasket may deform after heating to 90 ° C. DO NOT reuse the cassette and gasket. Discard them after this step. Use a new cassette and a new gasket in the subsequent steps.***

d. Add 400 μL 1X SSC (with 5% RI) to the new Stereo-seq Slide Cassette, incubate for 3 min then discard;

e. Repeat d for twice;

f. Prepare the blocking solution according to Table 4-3 below. The amount of blocking solution to prepare is 400 μL per slide, with 200 μL used for blocking and the other 200 μL used for preparing the antibody incubation solution. Add

200 μL /per slide of blocking solution to Stereo-seq Slide Cassette, make sure that the blocking solution cover the whole section and incubate for 20 minutes at room temperature;

Table 4-3 Blocking solution (one slide)

| Reagent | 1X (μL) |
|---|---|
| 1X SSC | 340 |
| Mixed serum | 40 |
| RI | 20 |
| Total | 400 |

g. While waiting for the blocking incubation to be done, prepare antibody solution according to Table 4-4, optimal dilution ratio is recommended by the manufacturer. After vertexing to mix well, briefly centrifuge, and then place on ice for use.

Table 4-4 Antibody solution (one slide)

| Reagent | 1X (μL) |
|---|---|
| Direct fluorescent antibody 1 | V1 |
| Direct fluorescent antibody 2<br>……<br>Direct fluorescent antibody n | V2<br>……<br>Vn |
| Blocking solution | 200 – (V1+V2+...+Vn) |
| Total | 200 |

h. Discard the blocking solution with a pipette. Slowly add the antibody solution from the non-tissue area until the solution covers the tissue section. Incubate at room temperature for 45 min in the dark.

***※ Make sure not to dry the tissue during the solution addition process. Otherwise, dried tissue might generate non-specific signals which interferes with the final results.***

***※ The fluorescent antibodies from Fluorophore 1 to Fluorophore n must be labeled with different fluorophores.***

##### 4.6 Imaging

The number of protein detections that can be achieved varies depending on the fluorescence channels available in different laboratory setups. For instance, if a laboratory's fluorescence microscope has seven fluorescence channels, it is possible to achieve twelve markers with thirteen colors (including DAPI) through two rounds of staining. If the laboratory only has four fluorescence channels, then through two rounds of staining, it is possible to achieve six markers with seven colors (including DAPI).

a. Precautions during the photography process.

1) Ensure that the chip is aligned with the edges of the imaging platform and do not tilt the chip;
2) Adjust the imaging parameters to ensure even exposure of the tissue area, with clear images and track lines;
3) It is recommended to use "manual focusing" for imaging. First, select model points at the four corners of the chip, then focus on several points both inside and outside the tissue to obtain clear track lines and tissue staining images simultaneously. Please use "auto focus" mode with caution. Auto focus may be difficult to adjust and may not be able to focus on both the track lines and the tissue at the same time;

4) When performing manual focusing for imaging, first select the appropriate focal plane, exposure, and illumination parameters to clearly identify the tissue contours. Then, adjust these parameters in the blank areas to clearly identify the Track lines. Avoid setting the photography parameters too low or too high, as this will affect image quality;
5) Do not touch the surfaces around the microscope during scanning. Avoid introducing any sources of vibration near the microscope, such as movement and walking;
6) Ensure that the elapsed time from tissue photography to imaging is within 30 min. Do not leave the chip in the same position for an extended period to avoid photobleaching of the fluorescence signal. Turn off the laser when the chip is not being imaged to prevent prolonged exposure, which could lead to local photobleaching;
7) Compressed or scaled images can affect image resolution and stitching results. Please ensure that the TIFF images exported from the microscope are neither compressed nor scaled to avoid image distortion that could impact downstream analysis;
8) After imaging, please immediately check if the images are clear and if the Track lines are distinct;
9) To minimize the impact on the transcriptome, it is recommended to take photos after achieving a clear focus to avoid multiple rounds of imaging;

b. Fluorescence photography operation steps;

This section provides a brief description of the general steps for fluorescence photography. For specific photography procedures, please refer to the operation manual of the fluorescence microscope configured in the laboratory.

1) Open the fluorescence microscope photography software, select the epi-fluorescence scanning mode, and create a new folder in a fluorescent

microscope-connected PC, and name it with the chip ID number and other essential information;

2) Turn on the fluorescence microscope and set the fluorescence channel;

3) Place 1 ~ 2 μL of water on the imaging platform first, then transfer and place the Stereo-seq Chip Slide onto the water drop. Water surface tension will grab onto the Slide and adhere it to the imaging platform;

4) Remove the light shield, switch to 4X objective lens,adjust brightness and gain. The specific parameters will vary depending on different microscopes. As long as the tissue can be imaged clearly, the light intensity should be kept in the lower range to prevent fluorescence quenching;

5) After scanning with 4X objective lens, take fluorescence images from the chip with the following microscope setting: select the fluorescence channel, 10X objective lens, full scan on capture area; As shown in Figure 1, taking the DAPI channel image of mouse brain tissue as an example, the red box in the middle picture is the selected tissue area, and the small blue boxes are the added focal points. The picture on the left shows a screenshot of a focal field window selected outside the tissue, which should ensure that the tracklines are clear and distinct. The picture on the right is a screenshot of a focal field window selected in the tissue area, which should ensure that the tissue outline and contour are clear;

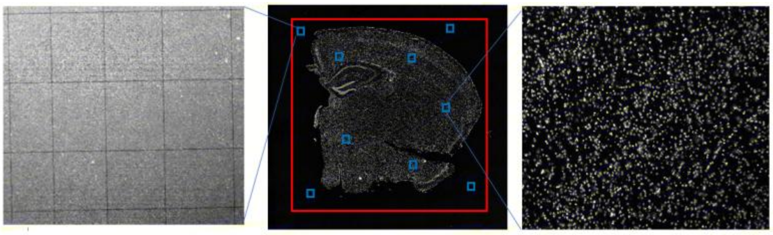

**Figure 1. DAPI image of mouse brain tissue**

6) Once complete the first scan of the fluorescence channel, switch to the next channel directly WITHOUT moving the Stereo-seq Chip Slide, re-scanning the map, re-adding the focal points, or modifying the red box of the selected

tissue area. Save original tile (FOV) image files and stitched images. Simply create a new folder, name it with the next antibody, and save the folder. Then adjust the focus and exposure until the stained tissue is clearly displayed. Finally, complete the full scan on the capturing area with 10X or 20X objective lens;

7) After the scanning is completed, continue imaging with the next channel until all the antibody IF images have been acquired;

8) Save original tile (FOV) image files and stitched images;

※ ***Note: The specific magnification for photography should be chosen based on the requirements of subsequent data analysis.***

c. Assessment of Photography Quality:

1) The image is clear, and the track lines are distinct:

① To ensure that both the tissue and the Track lines are in focus, select multiple focus points during imaging;

② Uniformly set the focus points based on the size and shape of the tissue, and establish 4 additional focus points at the four corners of the chip outside the tissue;

③ After imaging, it may be necessary to adjust the contrast and brightness to make the Track lines clearly visible;

2) No stitching errors;

3) Avoid photobleaching;

4) The tissue area should not exceed 80% of the chip area;

d. The criteria for clear imaging and distinct track lines can be referenced from the following images;

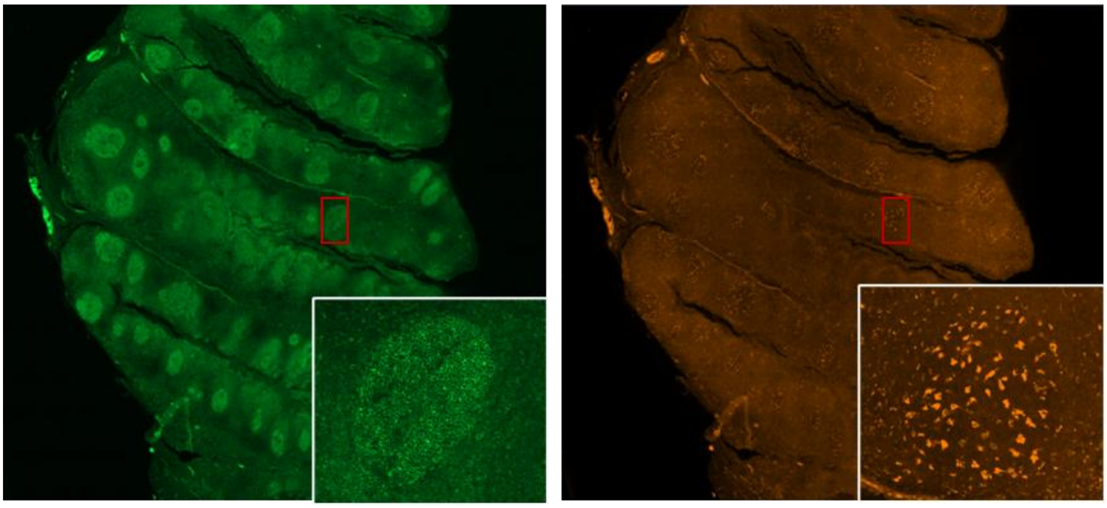

**Figure 2 Example of clear fluorescence signal**

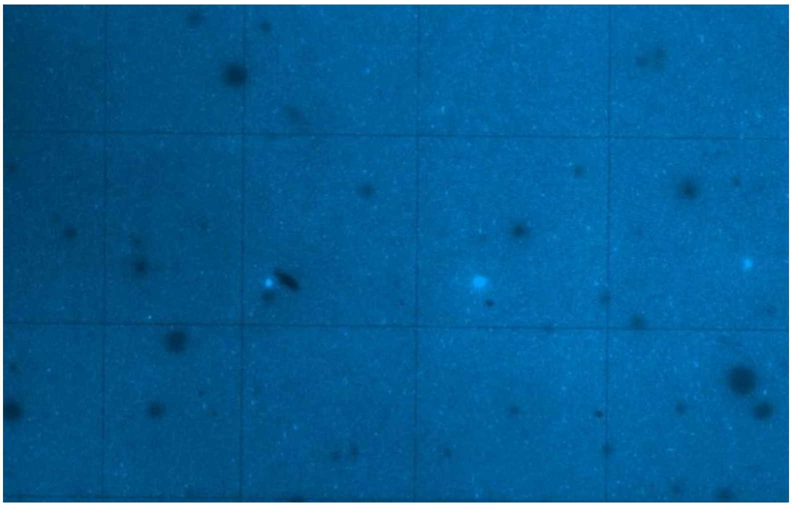

**Figure 3 Qualified image of clear tracklines**

#### 4.7 Antibody Elution

a. During fluorescence photography, place the PCR Adaptor in the PCR thermal cycler and set the program as follows:

Table 4-5 Antibody Elution program

| Temperature | Time |
|---|---|
| (Heated lid) 55°C | on |
| 50°C | ∞ |

b. Take out the antibody elution solution from 4°C 10 min in advance and place it in an EP tube on the PCR instrument heating module to preheat, 400 μL/chip;

c. After fluorescent imaging, immerse the coverslip-mounted Stereo-seq Chip Slide in a staining jar (alternatively, use a slide container or a 50 mL centrifuge tube) filled with at least 30 mL of 1X SSC, allow the coverslip to naturally detach from the tissue mounted chip, incubate for 3 min and gently rinse the Stereo-seq Chip Slide up and down to clean the chip surface;

d. Take out the Stereo-seq Chip Slide and wipe off the excess solution from around and the back of the slide with dust-free paper without touching the chip. Make sure there is no liquid residue around the chip;

e. Assemble the Cassette and Gasket then place the Stereo-seq Chip Slide in the Cassette according to the guide written in Appendix I: Stereo-seq Slide Cassette Assembly. It is recommended that you practice with a regular blank glass slide. Grip along the Stereo-seq Cassette to make sure the Stereo-seq Chip Slide has been locked in place;

f. Add 400 μL antibody elution buffer into the well of the Stereo-seq Slide Cassette, one droplet at each corner of the chip and then adding the rest of the solution to the middle to merge all the droplets. Apply sealing tape to the Stereo-seq Slide Cassette and make sure it is sealed tightly;

g. Place the Stereo-seq Slide Cassette in the 50°C PCR thermal cycler and let the chip incubate inside the PCR thermal cycler at 50°C for 20 min;

h. While waiting for the antibody elution process to be done, prepare antibody solution according to Table 4-6, optimal dilution ratio is recommended by the manufacturer. After vortexing to mix well, briefly centrifuge, and then place on ice for use;

Table 4-6 Second round antibody solution (one slide)

| Reagent | 1X (μL) |
|---|---|
| Direct fluorescent antibody 1 | V1 |
| Direct fluorescent antibody 2<br>……<br>Direct fluorescent antibody n | V2<br>……<br>Vn |
| Blocking solution | 200 – (V1+V2+...+Vn) |
| Total | 200 |

i. When decrosslinking is completed, carefully transfer the Stereo-seq Slide Cassette to the nearest bench and peel off the sealing tape. Immediately use a pipette to aspirate the antibody elution solution from the well, gently blow against the well walls several times, and then discard the antibody elution buffer with a pipette;

j. Add 400 μL 1X SSC (with 5% RI) to the Stereo-seq Slide Cassette, incubate for 3 min then gently blow against the well walls several times before discarding;

k. Repeat j for twice;

l. Add 200 μL of DMSO per chip from the corner of each well, Incubate for 10 min at room temperature;

m. Add 400 μL of 1X SSC (with 5% RI) per chip, gently pipe for several times.

##### 4.8 Second round of antibody incubation

a. Tilt the Stereo-seq Slide Cassette to as much as possible remove the 0.1 SSC (with 5% RI) from the chip surface, and then add the 200 μL pre-prepared second round of antibody incubation buffer onto the chip by first pipetting one droplet at each corner of the chip and then adding the rest of the solution to the middle to merge all the droplets. Incubate at room temperature for 45 min in the dark;

※ ***Note: This step does not require re-blocking; you can proceed directly to antibody incubation.***

b. While waiting for the antibody incubation to be done, prepare DAPI staining solution according to Table 4-7. Mix with pipette then leave it on ice in the dark until use;

c. Discard the antibody incubation buffer with a pipette. Add 200 μL/chip of 1X SSC (with 5% RI) incubate for 3 min then discard the liquid;

d. Repeat b for twice;

Table 4-7 DAPI staining solution (one slide)

| Reagent | 1X (μL) |
|---|---|
| 5X SSC | 60 |
| 50X-diluted DAPI solution | 2 |
| Nuclease-free water | 38 |
| Total | 100 |

##### 4.9 DAPI Staining and Fluorescence Imaging

a. Slowly add 200 μL/slide of DAPI staining solution to the tissue. Incubate for 2 min at room temperature in the dark;

b. Discard the DAPI staining solution with a pipette. Wash by adding 200 μL/slide of 1X SSC (5% RI) then discard the liquid;

c. Remove the slide from the Stereo-seq Slide Cassette and discard the cassette and gasket. Air dry the Stereo-seq Chip Slide or dry with a hand-held fan until no residual solution is left on the tissue;

d. Pipette 5 μL glycerol/slide gently onto the center of the tissue without introducing air bubbles. With a pair of forceps, place one end of the coverslip onto the glycerol covered tissue while holding the other end and then gradually lower the coverslip. Proceed to imaging immediately;

e. For fluorescence photography and image assessment, please refer to section 4.6;

##### 4.10 Decrosslinking

Equilibrate FFPE Decrosslinking Reagent to room temperature in advance;

a. After fluorescent imaging, immerse the coverslip-mounted Stereo-seq Chip Slide in a staining jar (alternatively, use a slide container or a 50 mL centrifuge tube) filled with at least 30 mL of 0.1X SSC, allow the coverslip to naturally detach from the tissue mounted chip, and gently rinse the Stereo-seq Chip Slide up and down to clean the chip surface;

b. Take out the Stereo-seq Chip Slide and wipe off the excess solution from around the back of the slide with dust-free paper without touching the chip. Make sure there is no liquid residue around the chip. The chip does not need to be air-dried;

c. Assemble the Cassette and Gasket then place the Stereo-seq Chip Slide in the Cassette according to the guide written in Appendix I: Stereo-seq Slide Cassette Assembly. It is recommended that you practice with a regular blank glass slide.

d. Grip along the Stereo-seq Cassette to make sure the Stereo-seq Chip Slide has been locked in place.

※ ***Make sure not to touch the front-side of the chip while assembling the Stereo- seq Slide Cassette.***

Place the PCR Adaptor in the PCR thermal cycler and set the program as follows:

Table 4-8 Decrosslinking incubation program

| Temperature | Time |
|---|---|
| (Heated lid) 85°C | on |
| 30°C | ∞ |
| 95°C | 25 min |
| 4°C | ∞ |

e. Add 400 μL FFPE Decrosslinking Reagent into the well of the Stereo-seq Slide Cassette. Apply sealing tape to the Stereo-seq Slide Cassette and make sure it is sealed tightly;

f. Place the Stereo-seq Slide Cassette on the PCR Adaptor in the PCR thermal cycler. Select Edit, and then select Next Step to skip the 30°C (Time = ∞) step;

g. Incubate at 95°C for 25 min. Meanwhile, prepare about 30 mL of methanol in a staining jar (alternatively, use a slide container or a 50 mL centrifuge tube), close the lid, and pre-cool it for 5 ~ 30 min at -20°C;

h. When decrosslinking is completed, carefully transfer the Stereo-seq Slide Cassette to the nearest bench and peel off the sealing tape. Discard the FFPE Decrosslinking Reagent with a pipette;;

i. Remove the slide from the Stereo-seq Slide Cassette according to instructions in Appendix I: Stereo-seq Slide Cassette Assembly, discard the cassette and gasket, then immediately proceed to the fixation step;

| Reagent | Purpose | Preparation |
|---|---|---|
| Methanol | Fixation | Pre-cooled for 5 ~ 30 min at –20˚C. |

#### 4.11 Fixation

a. Equilibrate the Stereo-seq Chip Slide to room temperature for 1 min, then immerse the tissue-mounted Stereo-seq Chip Slide in the pre-cooled methanol for a 20 min fixation at -20˚C. Make sure that the entire tissue section is completely submerged;

b. After fixation is completed, move the container to a sterile fume hood;

c. Take out the Stereo-seq Chip Slide and wipe off excess methanol from around and the back of the slide with dust-free paper without touching the chips. Make sure there is no methanol residue between chips;

d. Place the Stereo-seq Chip Slide on a slide staining rack and leave it in the fume hood for 4-6 min to let the methanol fully evaporate;

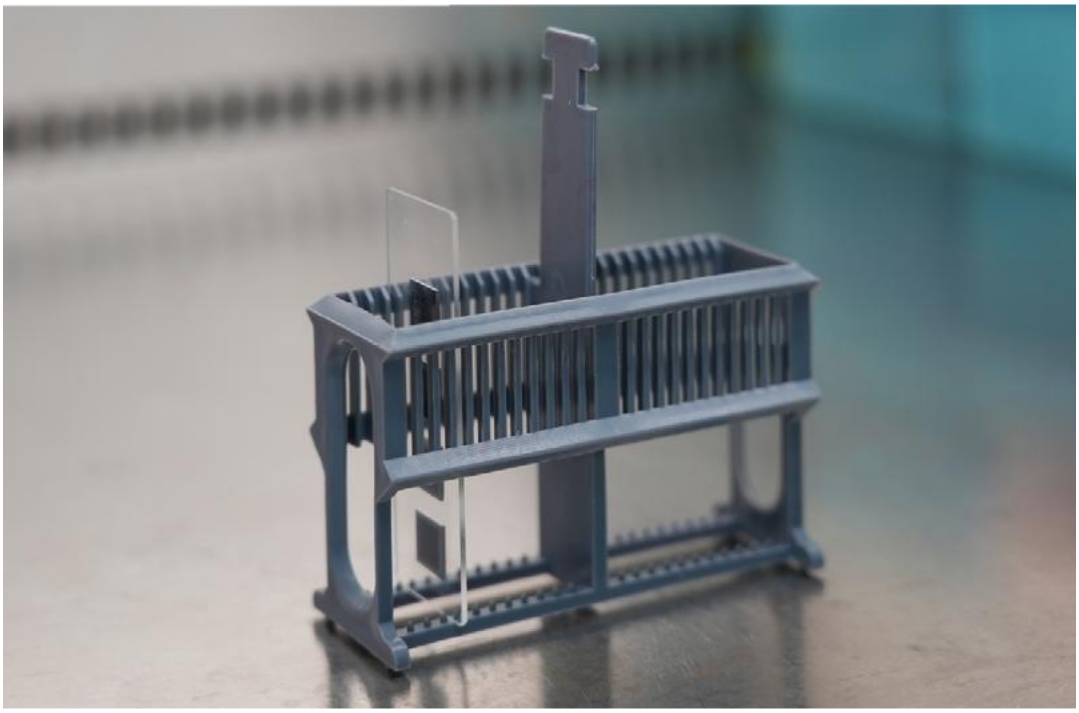

e. When the methanol is fully evaporated, transfer the Stereo-seq Chip Slide onto a flat and clean bench;

f. Assemble the Stereo-seq Slide Cassette with a new cassette and gasket;

g. Grip along the Stereo-seq Cassette to make sure the Stereo-seq Chip Slide has been locked in place.

**Do not touch the front of the chip while assembling the Stereo-seq Slide Cassette**

#### 4.12 Tissue Permeabilization

**Experiment Preparation**

| Reagent | Purpose | Preparation |
|---|---|---|
| 0.01N HCl | Permeabilization | Prepare at least 1 mL of 0.01 N HCl per sample. Configure HCl to 0.01N. Measure and make sure the pH = 2. |
| **Always use freshly prepared 0.01N HCl (pH = 2.0 ± 0.1)**. For pre-made 0.1N HCl and newly purchased HCl, check the pH prior to the experiments. | | |
| 10X Permeabilization Reagent Stock Solution | Permeabilization | Add 1 mL of freshly prepared 0.01 N HCl to dissolve PR Enzyme, and thoroughly mix the reagent by pipetting. |
| **Do not vortex the permeabilization enzyme. Mix by pipette before using. Aliquot this 10X stock solution to avoid freeze-thaw cycles.** | | |
| 1X Permeabilization Reagent Solution | Permeabilization | Make 1X PR solution (200 μL / chip) by diluting 10X PR stock solution with 0.01 N HCl. |
| 0.1X SSC | Dilution | Add 10 μL 20X SSC into 1990 μL nuclease-free water and mix well. |

a. Set aside the 2 mL of 0.01 N HCl and 1X Permeabilization Reagent Solution you prepared in 4.10 Experiment Preparation;

b. Make sure your PCR thermal cycler has been switched on and set to 37°C with the heated lid set to 42°C. Place the PCR Adaptor in the PCR thermal cycler and set the program as follows:

Table 4-9 Tissue permeabilization incubation program

| Temperature | Time |
|---|---|
| (Heated lid) 42°C | on |
| 37°C | ∞ |
| 37°C | 30 min |
| 4°C | ∞ |

c. Add 200 μL of 1X Permeabilization Reagent Solution onto the chip by first pipetting one droplet at each corner of the chip and then adding the rest of the solution to the middle to merge all the droplets. Apply sealing tape to the Stereo-seq Slide Cassette and make sure it is sealed tightly.

**Make sure the chip is completely covered with 1X Permeabilization Reagent Solution.**

d. Thaw FFPE RT Oligo and FFPE Dimer on ice.

e. Place the Stereo-seq Slide Cassette in the 37°C PCR thermal cycler and let the chip incubate inside the PCR thermal cycler at 37°C for 30 min.

f. While waiting for permeabilization to be completed, prepare FFPE MIX Solution according to below Table and leave it on ice until use. **【PREPARE AHEAD】**

Table 4-10 FFPE MIX Solution

| Components | 1X ( μL) | 2X + 10% ( μL) | 3X + 10% ( μL) | 4X + 10% ( μL) |
|---|---|---|---|---|
| **FFPE RT Buffer Mix** | 158 | 347.6 | 521.4 | 695.2 |
| **FFPE RT Enzyme Mix** | 30 | 66 | 99 | 132 |

| FFPE RT Oligo | 10 | 22 | 33 | 44 |
|---|---|---|---|---|
| **FFPE Dimer** | 2 | 4.4 | 6.6 | 8.8 |
| **Total** | 200 | 440 | 660 | 880 |

g. When incubation is completed, remove the Stereo-seq Slide Cassette from the PCR Adaptor (37°C);

h. Remove 1X Permeabilization Reagent Solution with a pipette from the corner of each well without touching the chip surface;

i. Add 200 μL of 0.1X SSC (with 5% RI) per chip from the corner of each well.

#### 4.13 FFPE MIX Reaction

a. Place the PCR Adaptor in the PCR thermal cycler in advance. Set the temperature to 42°C and the heated lid to 45°C according to the following program:

Table 4-11 FFPE MIX hybridization incubation program

| Temperature | Time |
|---|---|
| **(Heated lid) 45° C** | on |
| **42° C** | ∞ |

b. Gently add 200 μL of FFPE MIX Solution per chip along the side of each well, ensuring that the well surface is uniformly covered with FFPE MIX Solution. Apply sealing tape to the Stereo-seq Slide Cassette and make sure it is sealed tightly;

c. Place the Stereo-seq Slide Cassette in the 42°C PCR thermal cycler and let the chip incubate inside the PCR thermal cycler at 42°C for 5 hr or longer (no longer than 24 hr).

#### 4.14 cDNA Release and Collection

**If white precipitates are visible in the buffer, dissolve them by heating the buffer**

**at 55 ° C again and equilibrate to room temperature before mixing.**

| **Prepare** | | |
|---|---|---|
| **Reagent** | Preparation Steps | Maintenance |
| **cDNA Release** | Heat the buffer for 5 min at 55°C to dissolve the precipitate. Equilibrate it to room temperature prior to use. | Room temperature |

a. Remove the Stereo-seq Slide Cassette from the PCR Adaptor, discard FFPE MIX Solution, then wash one time with 200 μL 0.1X SSC per chip.

b. Prepare the cDNA Release Mix according to Table 4-12 and maintain the mix at room temperature.

Table 4-12 cDNA Release Mix

| **Components** | **1X ( μL)** | **2X + 10% ( μL)** | **3X + 10% ( μL)** | **4X + 10% ( μL)** |
|---|---|---|---|---|
| **cDNA Release Buffer** | 380 | 836 | 1254 | 1672 |
| **Enzyme** | 20 | 44 | 66 | 88 |
| **Total** | 400 | 880 | 1320 | 1760 |

c. Place the PCR Adaptor in the PCR thermal cycler in advance. Set the temperature to 55°C and the heated lid to 60°C according to the following Table 4-13 program:

Table 4-13 cDNA Release Incubation Program

| **Temperature** | **Time** |
|---|---|
| (Heated lid) 60°C | on |
| 55°C | 5 hr ~ 24 hr |
| 55°C | ∞ |

d. Add 400 μL of cDNA Release Mix per chip into each well of the Stereo-seq Slide Cassette;

e. Apply sealing tape to the Stereo-seq Slide Cassette and make sure it is sealed tightly. Incubate the Stereo-seq Slide Cassette at 55°C for 5 hr or longer (no longer than 24 hr);

**Stop Point:**

**cDNA collection step may be left overnight. If it is left overnight, make sure the Stereoseq Slide Cassette is sealed tightly with the sealing tape.**

f. Before collecting the eluted cDNA, pre-heat 500 μL of nuclease-free water per chip at 55°C for ≥ 30 min;

g. After the reaction, completely collect the cDNA Release Mix from each well into a new 2 mL centrifuge tube;

h. Add 350 μL of pre-heated nuclease-free water per chip into each well. Pipette up and down to wash the chip surface thoroughly and then collect it into the same 2 mL centrifuge tube with the cDNA Release Mix.

Collect as much volume as possible to retrieve enough cDNA from the chip. cDNA Release Mix should be about 400 μL after incubation (the volume might be less than 400 μL). You must combine the collected cDNA Release Mix with the 350 μL nuclease-free water before proceeding to the next step.

The Stereo-seq Chip Slide may be discarded. Ensure that all of the chip ID numbers have been recorded as required for downstream analysis.

#### 4.14 cDNA Purification and Amplification

##### Background Information

For bead-based purification, we recommend using DNA Cleanup Beads AMPure® XP (Agencourt, Cat. No. A63882), SPRIselect (Beckman Coulter, Cat. No. B23317/B23318/B23319) or VAHTS™ DNA Clean Beads (Vazyme, Cat. No. N411-02). If magnetic beads from other sources are used, please optimize the cleanup conditions before getting started.

##### Before Use

- To ensure the DNA capture efficiency of the magnetic beads, equilibrate the beads to room temperature 30 min before use;
- Vortex or pipette up and down to ensure that the beads are thoroughly mixed every time before use;
- The number of magnetic beads directly affects the distribution of purified DNA fragments.

##### Operation Notes

a. In the magnetic separation step, allow the solution to become completely clear before removing the supernatant. This process usually takes approximately 2 ~ 3 min, but can be longer or shorter depending on the type of magnetic separation rack being used.

b. When collecting the supernatant with a pipette after magnetic separation, avoid taking up the beads. Instead of collecting the entire supernatant fraction, leave 2 ~ 3 μL in the tube to avoid the pipette from direct contacting the beads. If the beads are mistakenly taken up, dispense everything and redo the magnetic separation.

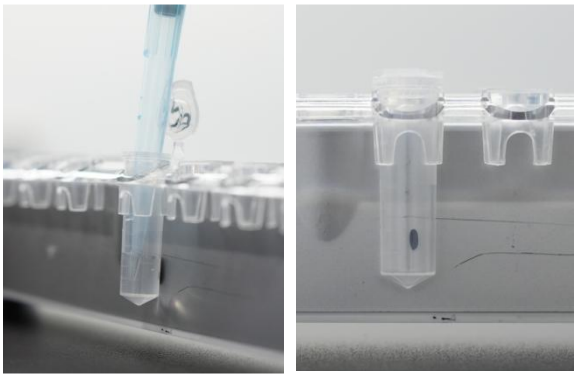

c. Use freshly prepared 80% ethanol (at room temperature) to wash the beads. Keep the sample tube on the magnetic separation rack during the washing step. Do not shake or disturb the beads

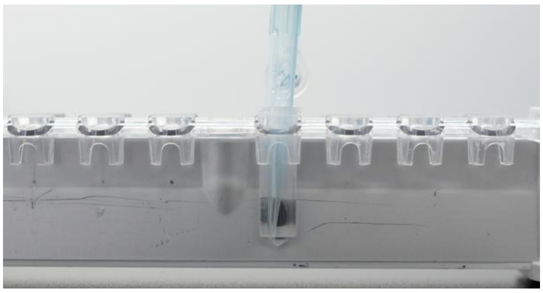

d. After the second washing of beads with ethanol, try to remove all of the liquid within the tube. You may centrifuge briefly to accumulate any remaining liquid at the bottom of the tube, then separate the beads magnetically, and remove the remaining liquid by using a small-volume pipette.

e. After washing twice with ethanol, air-dry the beads at room temperature. Drying usually takes approximately 5 ~ 10 min, depending on the lab temperature and humidity level. Watch closely until the pellet appears sufficiently dry with a matte appearance, then continue to the elution step with TE Buffer.

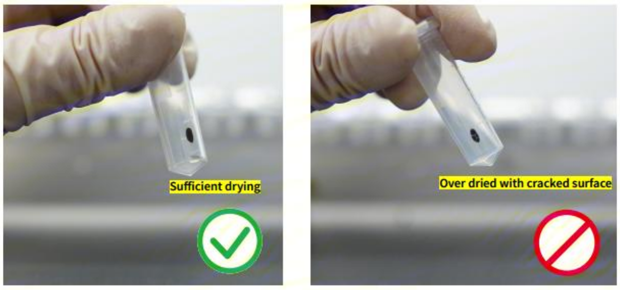


f. During the elution step, do not touch the beads with the pipette tip when removing the supernatant. Contamination of a DNA sample with beads may affect subsequent purification steps. Therefore, to prevent the pipette tip from directly contacting the beads, always collect the eluate in 2 μL less than the initial volume of TE Buffer used for the elution.

g. Pay attention when opening/closing the lid of a sample tube on a separation rack. Strong vibrations may cause samples or beads to spill from the tubes. Hold the body of the tube while opening the lid.

h. If white precipitates are visible in the collected cDNA, dissolve them by heating at 55˚C and equilibrate to room temperature before proceeding to the purification step. Equilibrate the magnetic beads to room temperature for at least 30 min.

i. cDNA Purification Procedures with 1X Magnetic Bead

1) Mix the collected cDNA (about 750 μL) with the beads in a ratio of 1 : 1. Vortex the mix then incubate it at room temperature for 10 min.

2) Spin down and place the tube onto a magnetic separation rack for 3 min until the liquid becomes clear.

3) Carefully remove and discard the supernatant with a pipette (if foam is visible on the cap, discard it with a pipette).

4) Keep the tube on the magnetic separation rack and add 1.5 mL of freshly prepared 80% ethanol. Wash the beads by rotating the tube on the magnetic rack. Incubate for 30 sec and carefully remove and discard the supernatant.

**Always place the pipette tips on the tube wall and away from the magnetic beads. Do not disturb the beads while transferring the supernatant (if foam is visible on the cap, clean the cap with 80% ethanol).**

5) Repeat step 4;

6) Keep the tube on the magnetic rack and open the lid to air-dry the beads at room temperature until no wetness (reflectiveness) or crack is visible. Drying times will vary but will take approximately 5 ~ 15 min;

7) Add 44 μL of nuclease-free water to the dried beads. Mix the beads and nucleasefree water by vortexing. Incubate at room temperature for 5 min. Spin down briefly and place the sample tube onto a magnetic separation rack for 3 ~ 5 min until the liquid becomes clear;

8) Transfer the supernatant (~ 42 μL cDNA) into a new 0.2 mL PCR tube;

j. If the volume of collected eluted cDNA is less than 42 μL, top it up with nuclease-free water.

Store the beads with 40 μL of nuclease-free water at 4°C after collecting the eluted cDNA untill your cDNA final product has passed QC.

k. Prepare PCR Mix according to Table 4-14. The total volume for the PCR reaction is 100 μL.

Table 4-14 PCR Mix

| Components | 1X ( μL) | X + 10% ( μL) | 3X + 10% ( μL) | 4X + 10% ( μL) |
|---|---|---|---|---|
| cDNA Amplification Mix | 50 | 110 | 165 | 220 |
| FFPE cDNA Primers Mix | 8 | 17.6 | 26.4 | 35.2 |
| Eluted cDNA | 42 | 2 x 42 | 3 x 42 | 4 x 42 |
| Total | 100 | 2 x 100 | 3 x 100 | 4 x 100 |

l. Mix gently and short spin before placing the reaction tube in a PCR thermal cycler.

m. Amplify the eluted cDNA based on the PCR program shown in Table 4-15.

Table 4-15 PCR program for amplification (for 100 μL)

| Temperature | Time | Cycle |
|---|---|---|
| (Heated lid) 105°C | on | / |
| 95°C | 5 min | 1 |
| 98°C | 20 s | 13 |
| 58°C | 20 s | |
| 72°C | 30 s | |
| 72°C | 5 min | 1 |
| 12°C | ∞ | / |
