## Supplementary 2 for "ST-FFPE-mIF: Integrating Spatial Transcriptomics and Multiplex Immunofluorescence in Formalin-Fixed Paraffin-Embedded Tissues Using Stereo-seq"

Table 2-1 Reagent preparation for tissue blocking and antibody incubation

| Reagent | Preparation Steps | Maintenance |
|---|---|---|
| Mixed serum | Retrieve aliquoted goat serum and horse serum from -20°C storage. Allow them to thaw and then mix them in 1:1 ratio in a tube. | On ice until use |
| Primary Antibodies | Take them out of -20°C or 4°C (depending on the manufacturer's instruction) and centrifuge at 14,000g, 4°C for 10 min then leave them on ice. | On ice until use |
| Secondary Antibodies | Take them out of -20°C or 4°C (depending on the manufacturer's instruction) and centrifuge at 14,000g, 4°C for 10 min then leave them on ice. | On ice until use |
| Diluted primary antibody (optional) | Primary antibodies can be diluted with blocking solution to a desired concentration if needed. | On ice until use |
| Diluted secondary antibody (optional) | Primary antibodies can be diluted with blocking solution to a desired concentration if needed. | On ice until use |
| DAPI | Take them out of -20°C or 4°C (depending on the manufacturer's instruction) and leave them on ice. | On ice until use |

d. Remove the 5-slide container from the water bath and allow it to cool down gradually at room temperature.

Notes:

- Tris-EDTA is a commonly used antigen retrieval method, but antigen retrieval can also be performed according to the specific requirements of each antibody.
- Antigen retrieval solution can be placed in containers other than the 5-slide container, depending on the situation.
- If using 5-slide containers as containers for antigen retrieval solution, the number of containers needed should be determined based on the number of slides.

e. Remove the first and last four sections and proceed with the routine immunohistochemistry (IHC) protocol.

#### 2.5 Tissue Blocking and Primary Antibody Incubation

a. After the antigen retrieval solution has cooled down, use forceps to remove the slides and place them into a staining jar that has been pre-filled with 1X SSC. Wash the slides on a shaker at room temperature incubate for 3 min and then discard. Repeat this step three times.

b. Take out the microscope slide and wipe off excess liquid from around and the back of the slide with dust-free paper without touching the tissue.

c. Use a Super PAP Pen (hydrophobic barrier pen) on the microscope slide to draw a circle around the tissue to create a hydrophobic exclusion zone to prevent the subsequent addition of fluid from escaping.

d. Prepare the blocking solution according to Table 2-2 below. The amount of blocking solution to prepare is 300 μL per slide, with 100 μL used for blocking, another 100 μL used for preparing the primary antibody incubation solution, the other 100 μL used for preparing the secondary antibody incubation solution. Add 100 μL per slide of blocking solution to hydrophobic exclusion zone surrounding the tissue. Adjust the volume according to the size of the hydrophobic circle (for example, if the circle is 0.5 cm X 0.5 cm, use 50 μL per slide), and incubate at room temperature for 20 minutes.

Table 2-2 blocking solution (one slide)

| Reagent | 1X (μL) |
|---|---|
| SSC | 270 |
| Mixed serum | 30 |
| Total | 300 |

e. While waiting for the incubation to be done, prepare the antibody solution according to Table 2-3. For the initial determination of antibody dosage, prepare the antibody solution according to the concentration recommended by the manufacturer. If the recommended dilution is 1:200, for example, add 0.5 μL of antibody. Mix the antibody incubation solution by pipetting, and then place it on ice for use.

Table 2-3 Primary antibody solution (one slide)

| Reagent | 1X (μL) |
|---|---|
| Primary antibody 1 | V1 |
| Primary antibody 2<br>......<br>Primary antibody n | V2<br>......<br>Vn |
| Blocking solution | 100 - (V1+V2+...+Vn) |
| Total | 100 |

f. Discard the blocking solution with a pipette. For experimental groups: Slowly add the antibody solution from the non-tissue area until the solution covers the tissue section. Do not exceed 100 μL/slide. Make sure to label the dilution ratio on the microscope slide. Incubate at room temperature for 45 min. For the negative control group: Add 100 μL/slide of blocking solution. Incubate at room temperature for 45 min.

g. While waiting for the primary antibody incubation to be done, prepare secondary antibody solution according to Table 2-4. After vortex mixing and brief centrifugation, leave the secondary antibody solution on ice in the dark until use.

h. Discard the primary antibody solution with a pipette. Wash by adding 100 μL/slide of 1X SSC. Incubate for 3 min and then discard. Repeat h twice for a total three-time wash;

i. For experimental groups: Slowly add the secondary antibody solution from the non-tissue area until the solution covers the tissue section. Do not exceed 100 μL/slide. Make sure to label the dilution ratio on the microscope slide. Incubate at room temperature for 25 min in the dark. For the negative control group: Add 100 μL/slide of blocking solution. Incubate at room temperature for 25 min.

Table 2-4 Secondary antibody solution (one slide)

| Reagent | 1X (μL) |
|---|---|
| Secondary antibody 1 | V1 |
| Secondary antibody 2<br>……<br>Secondary antibody n | V2<br>……<br>Vn |
| Blocking solution | 100 – (V1+V2+...+Vn) |
| Total | 100 |

※ ***Make sure not to dry the tissue during the solution addition process. Otherwise, dried tissue might generate non-specific signals which interferes with the final results.***

e. Repeat d for twice;

f. Prepare the blocking solution according to Table 4-3 below. The amount of blocking solution to prepare is 600 μL per slide, with 200 μL used for blocking, another 200 μL used for preparing the primary antibody incubation solution and the other 200 μL used for preparing the secondary antibody incubation solution. Add 200 μL per slide of blocking solution to Stereo-seq Slide Cassette, make sure that the blocking solution cover the whole section and incubate for 20 minutes at room temperature;

Table 4-3 blocking solution (one slide)

| Reagent | 1X (μL) |
|---|---|
| 1X SSC | 510 |
| Mixed serum | 60 |
| RI | 30 |
| Total | 600 |

g. While waiting for the blocking incubation to be done, prepare antibody solution according to Table 4-4, optimal dilution ratio is recommended by the manufacturer. After vertexing to mix well, briefly centrifuge, and then place on ice for use.

Table 4-4 Primary antibody solution (one slide)

| Reagent | 1X (μL) |
|---|---|
| Primary antibody 1 | V1 |
| Primary antibody 2<br>......<br>Primary antibody n | V2<br>......<br>Vn |
| Blocking solution | 200 – (V1+V2+...+Vn) |

| | |
|---|---|
| Total | 200 |

h. Discard the blocking solution with a pipette. Slowly add the antibody solution from the non-tissue area until the solution covers the tissue section. Incubate at room temperature for 45 min.

i. While waiting for the primary antibody incubation to be done, prepare secondary antibody solution in Table 4-5 according to the recommended dilution or manufacturer's instruction for each secondary antibody. After vortex mixing and brief centrifugation, leave the secondary antibody solution on ice in the dark until use.

Table 4-5 Primary antibody solution (one slide)

| Reagent | 1X (μL) |
|---|---|
| Secondary antibody 1 | V1 |
| Secondary antibody 2<br>......<br>Secondary antibody n | V2<br>......<br>Vn |
| Blocking solution | 200 – (V1+V2+...+Vn) |
| Total | 200 |

***※ Make sure not to dry the tissue during the solution addition process. Otherwise, dried tissue might generate non-specific signals which interferes with the final results.***

***※ The fluorescent antibodies from Fluorophore 1 to Fluorophore n must be labeled with different fluorophores.***

#### 4.6 Secondary Antibody Incubation

a. Discard the primary antibody solution with a pipette;

b. Wash by adding 400 μL/slide of 1X SSC. Incubate for 3 min then discard;

c. Repeat b. twice for a total three-time wash;

d. Slowly add 200 μL/slide of secondary antibody solution to the tissue. Incubate for 25 min at room temperature in the dark;

e. Discard the antibody solution with a pipette. Wash by adding 200 μL/slide of 1X SSC (with 5% RI) into the Stereo-seq Slide Cassette. Incubate for 3 min then discard. Repeat e twice for a total three-time wash;

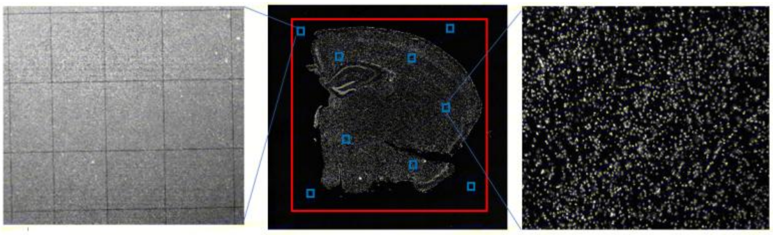

**Figure 1. DAPI image of mouse brain tissue**

6) Once complete the first scan of the fluorescence channel, switch to the next channel directly WITHOUT moving the Stereo-seq Chip Slide, re-scanning the map, re-adding the focal points, or modifying the red box of the selected tissue area. Save original tile (FOV) image files and stitched images. Simply create a new folder, name it with the next antibody, and save the folder. Then adjust the focus and exposure until the stained tissue is clearly displayed. Finally, complete the full scan on the capturing area with 10X or 20X objective lens;

7) After the scanning is completed, continue imaging with the next channel until all the antibody IF images have been acquired;

8) Save original tile (FOV) image files and stitched images;

※ ***Note: The specific magnification for photography should be chosen based on the requirements of subsequent data analysis.***

c. Assessment of Photography Quality

1) The image is clear, and the track lines are distinct:

① To ensure that both the tissue and the Track lines are in focus, select multiple focus points during imaging;

② Uniformly set the focus points based on the size and shape of the tissue, and establish 4 additional focus points at the four corners of the chip outside the tissue;

③ After imaging, it may be necessary to adjust the contrast and brightness to make the Track lines clearly visible;

2) No stitching errors;

3) Avoid photobleaching;

4) The tissue area should not exceed 80% of the chip area;

d. The criteria for clear imaging and distinct track lines can be referenced from the following images;

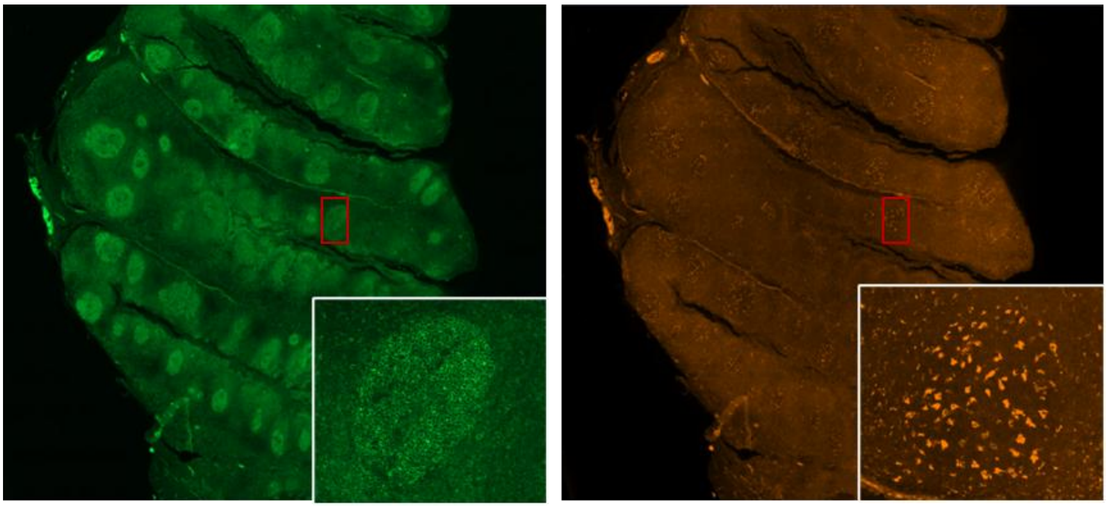

**Figure 2 Example of clear fluorescence signal**

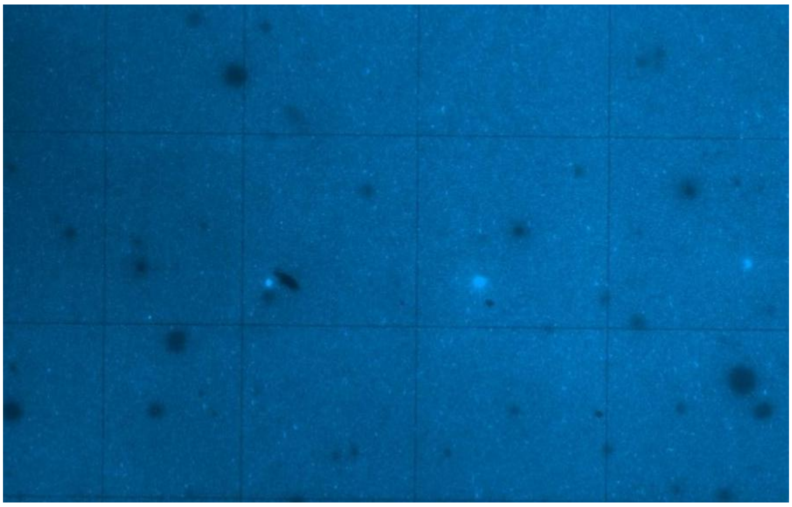

**Figure 3 Qualified image of clear tracklines**

### 4.8 Antibody Elution

a. During fluorescence photography, place the PCR Adaptor in the PCR thermal cycler and set the program as follows:

Table 4-7 second round primary antibody solution (one slide)

| Reagent | 1X (μL) |
|---|---|
| Primary antibody 1 | V1 |
| Primary antibody 2<br>……<br>Primary antibody n | V2<br>……<br>Vn |
| Blocking solution | 200 – (V1+V2+…+Vn) |
| Total | 200 |

i. When antibody elution is completed, carefully transfer the Stereo-seq Slide Cassette to the nearest bench and peel off the sealing tape. Immediately use a pipette to aspirate the antibody elution solution from the well, gently blow against the well walls several times, and then discard the antibody elution buffer with a pipette;

j. Add 400 μl 1X SSC (with 5% RI) to the Stereo-seq Slide Cassette, incubate for 3 min then gently blow against the well walls several times before discarding;

k. Repeat j for twice ;

l. Add 200 μL of DMSO per chip from the corner of each well, Incubate for 10 min at room temperature.

m. Add 400 μL of 1X SSC (with 5% RI) per chip, gently pipe for several times.

### 4.9 Second Round of Antibody Incubation

a. Tilt the Stereo-seq Slide Cassette to as much as possible remove the 0.1X SSC (with 5% RI) from the chip surface, and then add the 200 μL pre-prepared second round of primary antibody incubation buffer onto the chip by first pipetting one droplet at each corner of the chip and then adding the rest of the solution to the middle to merge all the droplets. Incubate at room temperature for 45 min;

b. While waiting for the primary antibody incubation to be done, prepare secondary antibody solution in Table 4-8 according to manufacturer's instruction for each secondary antibody. After vortex mixing and brief centrifugation, leave the secondary antibody solution on ice in the dark until use.

※ ***Note: This step does not require re-blocking; you can proceed directly to antibody incubation.***

c. Discard the primary antibody solution with a pipette;

Table 4-8 second round secondary antibody solution (one slide)

| Reagent | 1X (μL) |
|---|---|
| Secondary antibody 1 | V1 |
| Secondary antibody 2<br>……<br>Secondary antibody n | V2<br>……<br>Vn |
| Blocking solution | 200 – (V1+V2+…+Vn) |
| Total | 200 |

d. Wash by adding 400 μL/slide of 1X SSC. Incubate for 3 min then discard;

e. Repeat d twice for a total three-time wash;

f. Slowly add 200 μL/slide of secondary antibody solution to the tissue. Incubate for 25 min at room temperature in the dark;

#### 4.10 DAPI Staining and Fluorescence Imaging

a. While waiting for the antibody incubation to be done, prepare DAPI staining solution according to Table 4-9. Mix with pipette then leave it on ice in the dark until use;

b. Discard the antibody solution with a pipette. Wash by adding 200 μL/slide of 1X SSC (with 5% RI) into the Stereo-seq Slide Cassette. Incubate for 3 min then discard.

c. Repeat b twice for a total three-time wash;

d. Slowly add 200 μL/slide of DAPI staining solution to the tissue. Incubate for 2 min at room temperature in the dark;

e. Discard the DAPI staining solution with a pipette. Wash by adding 200 μL/slide of 1X SSC (with 5% RI) then discard the liquid;

f. Remove the slide from the Stereo-seq Slide Cassette and discard the cassette and gasket. Air dry the Stereo-seq Chip Slide or dry with a hand-held fan until no residual solution is left on the tissue;

g. Pipette 5 μL glycerol/slide gently onto the center of the tissue without introducing air bubbles. With a pair of forceps, place one end of the coverslip onto the glycerol covered tissue while holding the other end and then gradually lower the coverslip. Proceed to imaging immediately;

h. For fluorescence photography and image assessment, please refer to section 4.7;

Table 4-9 DAPI staining solution (one slide)

| Reagent | 1X (μL) |
| --- | --- |
| 5X SSC | 60 |
| 50X-diluted DAPI solution | 2 |
| Nuclease-free water | 38 |
| Total | 100 |

d. Place the Stereo-seq Chip Slide on a slide staining rack and leave it in the fume hood for 4-6 min to let the methanol fully evaporate;

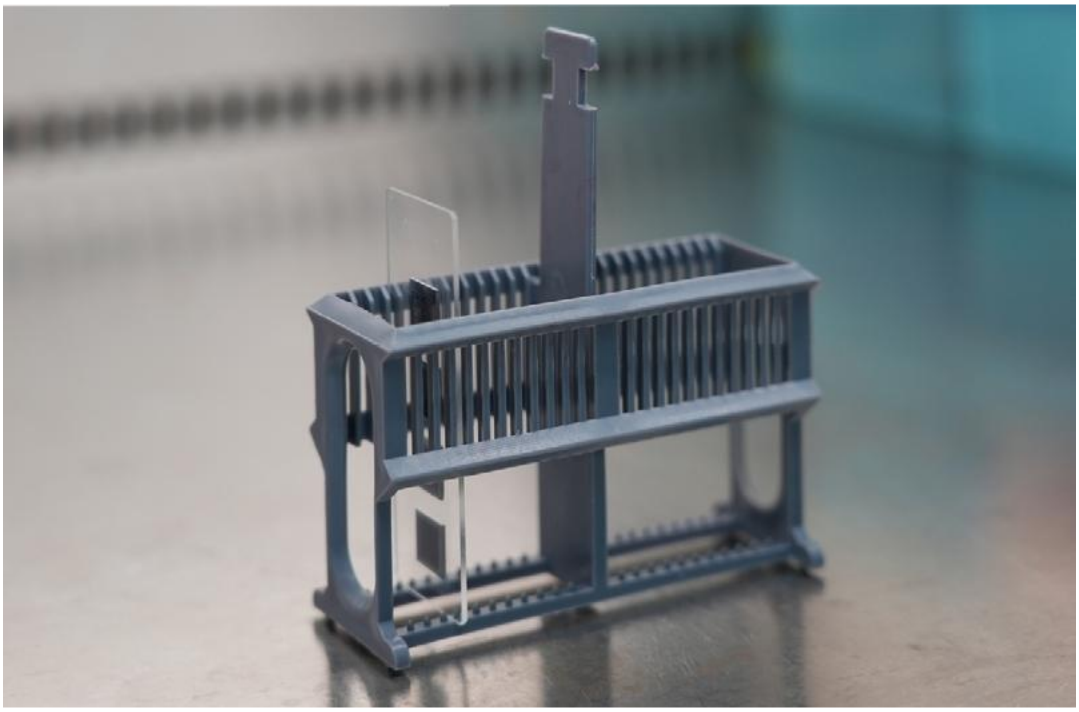

e. When the methanol is fully evaporated, transfer the Stereo-seq Chip Slide onto a flat and clean bench;

| | |
|---|---|
| | Equilibrate it to room temperature prior to use. |

a. Remove the Stereo-seq Slide Cassette from the PCR Adaptor, discard FFPE MIX Solution, then wash one time with 200 μL 0.1X SSC per chip.

b. Prepare the cDNA Release Mix according to Table 4-14 and maintain the mix at room temperature.

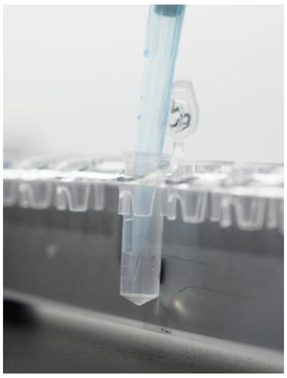

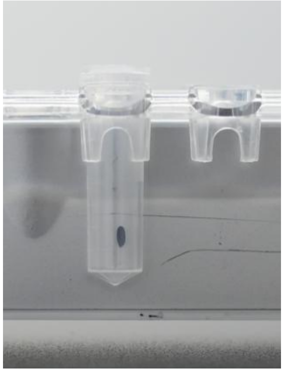

c. Use freshly prepared 80% ethanol (at room temperature) to wash the beads. Keep the sample tube on the magnetic separation rack during the washing step. Do not shake or disturb the beads

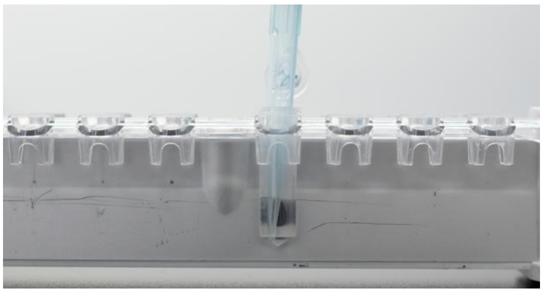

d. After the second washing of beads with ethanol, try to remove all of the liquid within the tube. You may centrifuge briefly to accumulate any remaining liquid at the bottom of the tube, then separate the beads magnetically, and remove the remaining liquid by using a small-volume pipette.

e. After washing twice with ethanol, air-dry the beads at room temperature. Drying usually takes approximately 5 ~ 10 min, depending on the lab temperature and humidity level. Watch closely until the pellet appears sufficiently dry with a matte appearance, then continue to the elution step with TE Buffer.

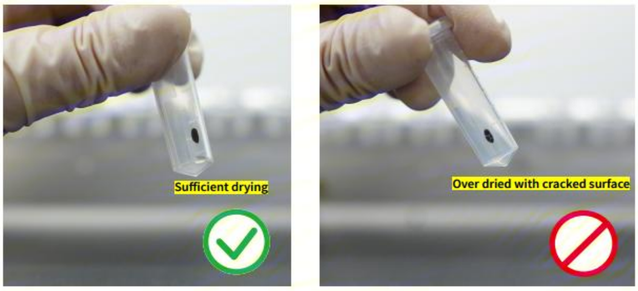

f. During the elution step, do not touch the beads with the pipette tip when removing the supernatant. Contamination of a DNA sample with beads may affect subsequent purification steps. Therefore, to prevent the pipette tip from directly contacting the beads, always collect the eluate in 2 μL less than the initial volume of TE Buffer used for the elution.

k. Prepare PCR Mix according to Table 4-16. The total volume for the PCR reaction is 100 μL.

Table 4-16 PCR Mix

| Components | 1X ( μL) | X + 10% ( μL) | 3X + 10% ( μL) | 4X + 10% ( μL) |
|---|---|---|---|---|
| cDNA Amplification Mix | 50 | 110 | 165 | 220 |
| FFPE cDNA Primers Mix | 8 | 17.6 | 26.4 | 35.2 |
| Eluted cDNA | 42 | 2 x 42 | 3 x 42 | 4 x 42 |
| Total | 100 | 2 x 100 | 3 x 100 | 4 x 100 |

l. Mix gently and short spin before placing the reaction tube in a PCR thermal cycler.

m. Amplify the eluted cDNA based on the PCR program shown in Table 4–17

Table 4–17 PCR program for amplification (for 100 μL)

| Temperature | Time | Cycle |
|---|---|---|
| (Heated lid) 105°C | on | / |
| 95°C | 5 min | 1 |
| 98°C | 20 s | 13 |
| 58°C | 20 s | |
| 72°C | 30 s | |
| 72°C | 5 min | 1 |
| 12°C | ∞ | / |
